# Characterization of altered molecular mechanisms in Parkinson’s disease through cell type-resolved multi-omics analyses

**DOI:** 10.1101/2022.02.13.479386

**Authors:** Andrew J. Lee, Changyoun Kim, Seongwan Park, Kyoungho Jun, Junghyun Eom, Seung-Jae Lee, Sun Ju Chung, Robert A. Rissman, Jongkyeong Chung, Eliezer Masliah, Inkyung Jung

**Author notes:** Correspondence to Eliezer Masliah and Inkyung Jung. These authors contributed equally to this work (Andrew J. Lee, Changyoun Kim).

## Abstract

Parkinson’s disease (PD) is a progressive neurodegenerative disorder. However, cell type-dependent transcriptional regulatory programs responsible for PD pathogenesis remain elusive. Here, we establish transcriptomic and epigenomic landscapes of the substantia nigra (SN) by profiling 87,733 nuclei obtained from healthy controls and PD patients. Our multi-omic data integration provides functional annotation of 128,724 *cis*-regulatory elements (*c*REs) and uncovers cell-type specific dysregulated *c*REs with a strong transcriptional influence on genes implicated in PD. The establishment of high-resolution three-dimensional chromatin contact maps identifies 656 target genes of dysregulated *c*REs and genetic risk loci, including both novel candidates and known PD risk genes. Notably, these new PD candidate genes exhibit modular gene expression patterns with unique molecular signatures in distinct cell types. Thus, our single-cell transcriptome and epigenome uncover cell type-specific disrupted transcriptional regulations in PD.

**Teaser:** Single-cell transcriptome and epigenome uncover cell type-specific disrupted transcriptional regulations in Parkinson’s disease.

## Introduction

Parkinson’s disease (PD) is a chronic, progressive neurodegenerative disorder accompanying both motor and non-motor symptoms including resting tremor, bradykinesia, rigidity, and other non-motor symptoms. Pathologically, PD is characterized by dopaminergic neuronal loss, abnormal protein deposition of Lewy bodies (LBs) and Lewy neurites (LNs), and neuroinflammation in the substantia nigra (SN) (*1–3*). With regard to its etiology, genetic studies for familial forms of PD, which accounts for ∼10% of PD patients, have identified critical causal genetic factors (*4–7*). Large-scale genome-wide association studies (GWAS) have further identified up to 90 genomic loci associated with PD (*8–13*). However, the identified genetic risk factors explain only 30% of familial and 3-5% of sporadic PD cases, ascribed to the complex genetic predisposition associated with the disease susceptibility (*13–16*). The interpretation of these genetic variants is hindered because specific cell types in which these genetic variants exert their function are unknown. Further, the functional mechanism by which these variants contribute to disease susceptibility is still elusive, as most of them are located in non-coding sequences.

The growing recognition that perturbations in regulatory elements involve in disease-specific gene expression and co-localize with many non-coding genetic variants provides a rationale for in-depth investigation of epigenome associated with PD (*17–21*). Although a systematic examination of *cis*-regulatory elements (*c*REs) in PD is scarce, a global dysregulation of lysine H3K27 acetylation (H3K27ac) landscape and a localization of PD GWAS genes proximal to the dysregulated *c*REs have been reported in the prefrontal cortex (*22*). Further, the advent of single-nucleus sequencing approach has allowed the investigation of epigenome landscape across individual brain cell types. A recent study reported the association of genetic variants of Alzheimer’s disease (AD) and PD at region- and cell type-specific *c*REs in the healthy brains (*23*). Another study has characterized AD-associated dysregulation in chromatin accessibility at sub-cell type level, identifying cell type-specific *c*RE candidates (*24*). In line with this, there is a tremendous demand for cell type-resolved investigation of aberrant *cis*-regulatory regions in PD with respect to PD-specific gene expression.

As the function of *c*REs is dependent on the long-range chromatin interaction with a target promoter located over large genomic distances, the identification of *c*RE-to-promoter relationships is challenging. To this end, high-throughput chromosome conformation capture (3C) methods, including ChIA-PET, Hi-C, capture Hi-C, HiChIP, and PLAC-seq, have allowed the investigation of genome-wide chromatin interactions (*25–29*), and substantially advanced our view on the functional role of *c*REs regulating distal target gene expression (*20, 30–33*). Recent integrative analyses of long-range chromatin interactome in healthy brain tissues have identified putative target genes of AD and PD genetic variants (*23, 34*). Nevertheless, the connection between *cis*-regulatory elements and altered gene expression in PD is largely unknown due to the lack of high-resolution three-dimensional (3D) chromatin contact maps available in the SN region of PD and control individuals.

To determine how cell type-dependent dysregulation in *cis*-regulatory regions affect molecular mechanisms related to PD pathogenesis in context of 3D chromatin interactions, we conducted an integrative analysis of multi-omic data generated from the SN, the brain region most affected by PD. Single-nucleus sequencing of RNA (snRNA-seq) and chromatin accessibility (snATAC-seq) established cell type-resolved transcriptome and epigenome for both PD and control SN. Analysis of global H3K27ac signals from chromatin immunoprecipitation followed by sequencing (ChIP-seq) identified a set of dysregulated *c*REs, which were annotated on the basis of active cell type. By integrating PD GWAS-variants, we confirmed a strong association between the cell-type resolved epigenome and the PD-associated genetic components. Furthermore, we generated high-resolution 3D chromatin contact maps to effectively expand potential PD candidate genes by inferring putative targets of dysregulated *c*REs and PD GWAS-SNPs. Notably, modular expression patterns of the putative target genes resolved heterogeneous molecular pathways involved in PD, in which we annotated each cluster with a unique biological property and responsible cell type. Our findings shed light on the complex molecular characteristics of PD pathogenesis, and expand candidate genes through mapping the target genes of PD-specific non-coding sequences in a cell type-resolved manner.

## Results

### Single-nucleus profiling of transcriptome and chromatin accessibility in the human SN of control and PD cases

To dissect the disease-specific gene regulations in PD, we conducted integrative multi-omics analysis on the transcriptome and chromatin accessibility for individual cell types present in the human SN (Fig. 1A). We performed snRNA-seq (10x Genomics v3) to characterize PD-associated changes in transcriptome at single-nucleus resolution in 15 flash-frozen postmortem SN specimens (late-stage PD = 4, control = 11) (table S1) (*35*). In parallel, we conducted snATAC-seq (10x Genomics v1) on 7 postmortem SN specimens (late-stage PD = 4, control = 3) and integrated 2 additional SN samples (*24*) to characterize cell type-resolved accessible chromatin (table S1). The SN specimens used for the experiments were obtained from Alzheimer’s Disease Research Center at the University of California, San Diego (fig. S1A). A rigorous quality control yielded a final set of 42,607 nuclei (30,534 control and 12,073 PD nuclei) for snRNA-seq and 45,126 nuclei (24,628 control and 20,498 PD nuclei) for snATAC-seq (Fig. 1, B and C).

**Fig. 1.**
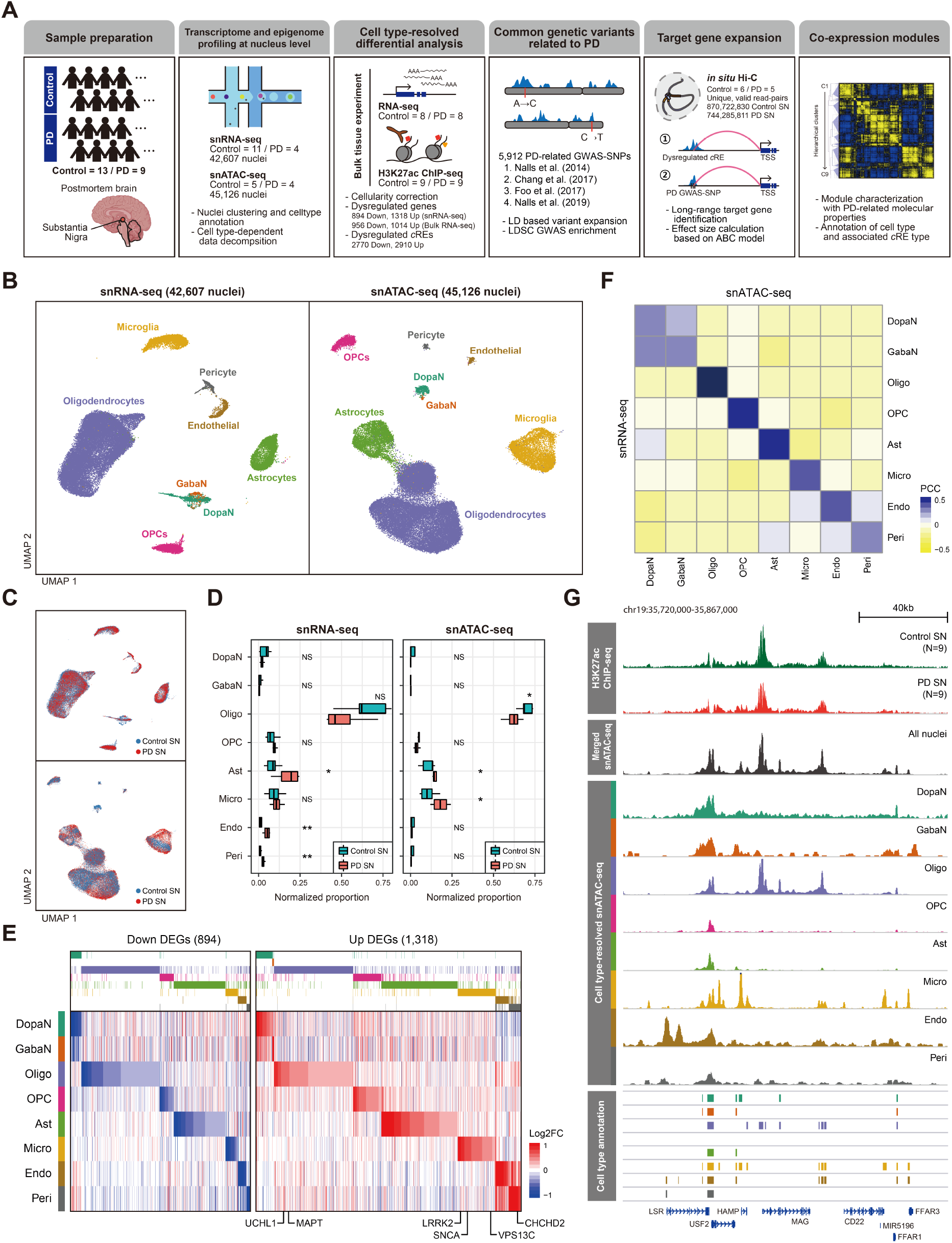
Single-nucleus profiling of transcriptomic and epigenomic landscape in the human SN. **(A)** A schematic of the study design, illustrating the preparation of sequencing-based omics data and downstream computational analysis. **(B)** UMAP embeddings of QC-passed nuclei for snRNA-seq (left) and snATAC-seq (right). The nuclei were annotated into dopaminergic neurons (DopaN), GABAergic neurons (GabaN), oligodendrocyte, oligodendrocyte precursor cells (OPCs), astrocytes (Ast), microglia (Micro), endothelial cells (Endo), and pericytes (Peri), based on cell type markers. **(C)** UMAP embeddings of nuclei colored by pathological status, where red and blue indicate nuclei from PD and control SN, respectively. **(D)** Box plots showing the proportion of nuclei mapped to each cell type and sample, compared between PD (red) and control (green) SN for snRNA-seq (left) and snATAC-seq (right). Two-sided Wilcoxon tests are performed to measure the statistical significance of the difference in cellular compositions between PD and control cases (*: p-value < 0.05, **: p-value < 0.01). The box boundaries and line correspond to the interquartile range (IQR) and median, respectively. Whiskers extend to the lowest or highest data points that are no further than 1.5 times the IQR from the box boundaries. **(E)** A heatmap by log2 fold change of cell type-resolved snRNA-seq reads (PD/control), illustrating 894 down-regulated DEGs (left) and 1,318 up-regulated DEGs (right), with the annotation of known PD risk genes. **(F)** A heatmap showing Pearson’s correlation coefficients (PCC) between snATAC-seq gene activity scores and snRNA-seq gene expression across the cell types present in the human SN. **(G)** Genome browser tracks of H3K27ac ChIP-seq signals for PD (red) and control (green) SN, and pseudobulk snATAC-seq signals resolved according to the cell types, along with tracks indicating the positions of cell type-resolved *c*REs. The signals for ChIP-seq and cell type-resolved snATAC-seq were normalized by the total reads mapped in *c*REs.

Unbiased clustering of the nuclei did not find a particular segregation by potential confounders (sex, age, read depth, and double score) in both snRNA-seq and snATAC-seq clusters (fig. S1, B and C, and fig. S2). On the basis of known marker genes, we profiled all major cell types in the SN, including neurons (*SYT1*), oligodendrocytes (Oligo; *MAG* and *MOBP*), oligodendrocyte precursor cells (OPCs; *PDGFRA*), astrocytes (Ast; *AQP4* and *GFAP*), microglia (Micro; *CD74* and *RUNX1*), endothelial cells (Endo; *CLDN5*), and pericytes (Peri; *PDGFRB*) (fig. S3A and fig. S4A). Sub-clustering of neuronal populations allowed the annotation of dopaminergic neurons (DopaN; *TH* and *SLC6A3*) as a distinct cluster from GABAergic neurons (GabaN; *GAD1* and *GAD2*) (fig. S3B and fig. S4B). Our cellular composition analysis indicated a moderate reduction of dopaminergic neurons (*P* = 0.24 for snRNA-seq, *P* = 0.11 for snATAC-seq) and a significant increase in astrocyte population (*P* = 0.038 for snRNA-seq, *P* = 0.049 for snATAC-seq), which may indicate that astrogliosis was accompanied as a neuroprotective mechanism in response to dopaminergic neuronal loss (Fig. 1D) (*36, 37*).

Further, we identified 2,212 cell type-specific differentially expressed genes (DEGs; down = 894, up = 1,318) (table S2) by iteratively performing a differential analysis in individual cell types based on single-nucleus transcriptome (Fig. 1E and fig. S5A). Interestingly, down-regulated DEGs were over-represented by cell type specific Gene Ontology (GO) pathways relevant to PD neuropathology, as exemplified by vesicle transport in oligodendrocytes, protein localization in astrocytes, and immune responses in microglia (fig. S5B). Enriched biological processes in up-regulated DEGs exhibited recurrent cellular processes in multiple cell types, including mitochondrial membrane organization, response to unfolded proteins, and regulation of protein ubiquitination (fig. S5C). We found that 6 of 20 well-known PD risk genes (*38*) were included in DEGs, which were concentrated in up-regulated DEGs with diverse cell type functionality, including *SNCA* (Oligo, Micro), *MAPT* (Oligo, OPC), *UCHL1* (Oligo), *LRRK2* (Micro), *VPS13C* (Micro), and *CHCHD2* (Peri) (Fig. 1E). Our results indicate the transcriptomic profiling at the single nuclei level effectively recapitulates PD biology.

We established cell type-resolved regulatory landscapes by decomposing the snATAC-seq reads according to cell type annotation. We confirmed that key marker genes obtained from snRNA-seq and snATAC-seq showed a high degree of overlap in all cell types (Fig. 1G, fig. S6, and table S3). In total, the 128,724 *c*REs were identified from pseudobulk snATAC-seq chromatin accessibility, which were assigned to the corresponding cell type (Fig. 1F and fig. S7A). A small fraction of *c*REs (1.12%) was commonly annotated by all cell types, which suggests that a high degree of cell type specificity was captured by the dynamic *c*RE repertoire (fig. S7B).

### Identification of dysregulated *cis*-regulatory elements in distinct cell types

Next, we questioned the presence of global dysregulation in the non-coding regulatory landscape associated with PD in individual cell types. While single-nucleus sequencing approach provides valuable insights into cell type specificity in the brain, it is challenging to effectively identify disease-specific dysregulated *c*REs due to its inherent sparsity. To compensate such limitations, we incorporated high-quality bulk histone H3 27th acetylation (H3K27ac) ChIP-seq results (late-stage PD = 9, control = 9) utilizing the SN specimens (fig. S7C). Given a high consistency between H3K27ac ChIP-seq and pseudobulk snATAC-seq signals across the *c*REs (Fig. 1G and fig. S7D), we implemented an iterative cellularity correction approach to resolve cellular heterogeneity in our bulk H3K27ac ChIP-seq results (fig. S8). Then, we performed a quasi-likelihood test in EdgeR (*39*) to identify 5,680 dysregulated *c*REs (down = 2,770, up = 2,910; Benjamini-Hochberg (BH)-adjusted *Q* < 0.05) (Fig. 2A and table S4). 0.93% of down-regulated *c*REs were found to be cell type common, retaining a similar level of cell type specificity as the overall *c*REs, with the cell types dominantly annotated in oligodendrocytes, OPCs, and astrocytes. A higher rate of cell type common *c*REs (5.01%) was found in up-regulated *c*REs, which may be ascribed to the recurrent cellular processes associated with up-regulated DEGs.

**Fig. 2.**
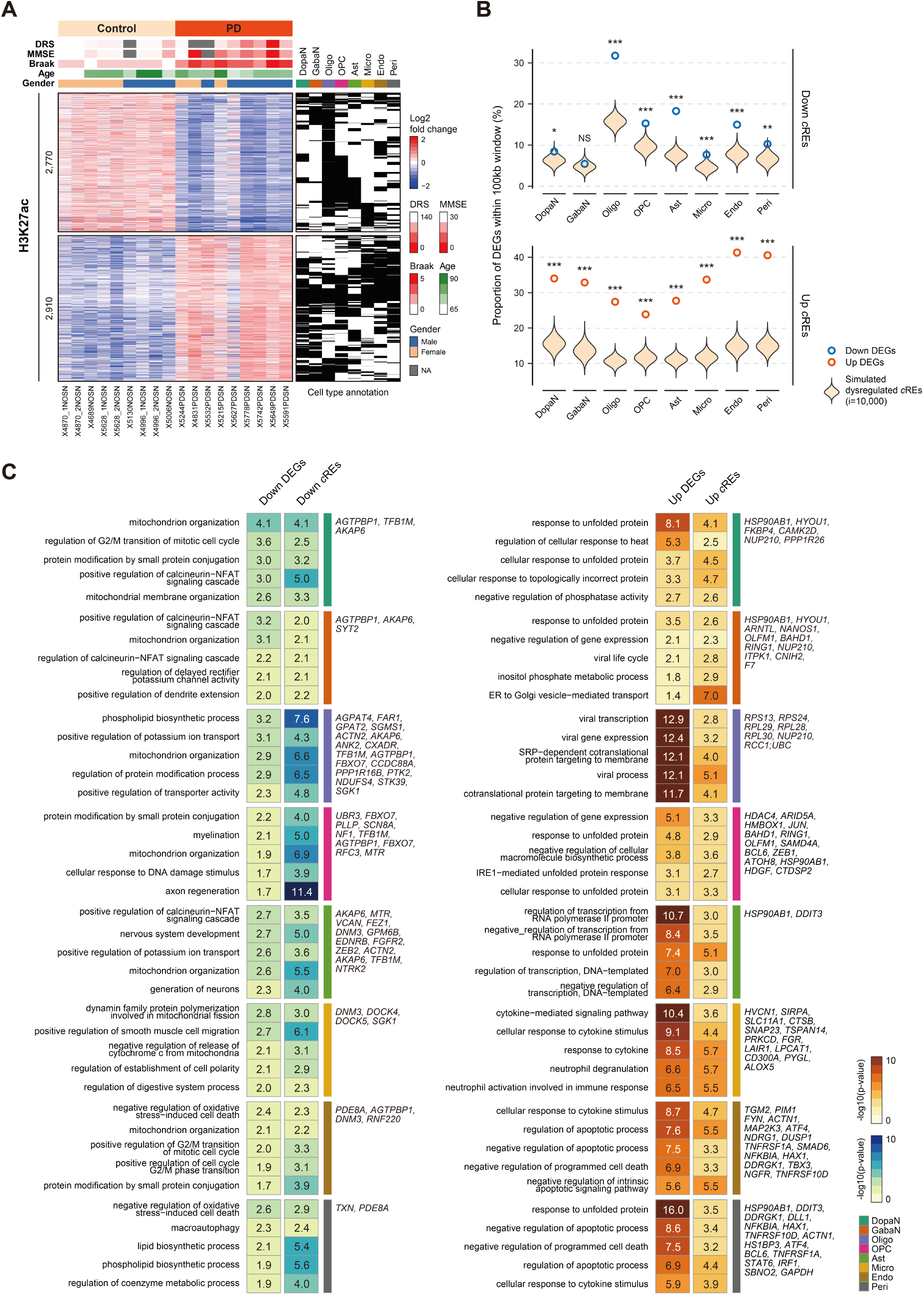
Dysregulation of *cis*-regulatory elements shaping PD-specific gene expression. **(A)** A heatmap of 2,770 down- and 2,910 up-regulated *c*REs by log2 fold change of normalized H3K27ac ChIP-seq reads divided by the mean of all samples, along with binarized annotation of cell types. **(B)** Enrichment analysis for colocalization between DEGs and dysregulated *c*REs in a distance genomic window (100kb). Violin plots represent the expected proportion of DEGs harbored by simulated *c*REs with 10,000 permutations. The observed proportion of DEGs shown in bold donuts (blue for down-regulated DEGs and dark orange for up-regulated DEGs). The statistical significance was calculated based on empirical testing (NS: not significant, *: p-value < 0.05, **: p-value < 0.01, ***: p-value < 0.001). **(C)** Top 5 enriched GO biological pathways for down- (left) and up- (right) regulated DEGs commonly represented by genes annotated by dysregulated *c*REs through Genomic Regions Enrichment of Annotations Tool (GREAT) for individual cell types. DEGs within PD-related pathways that are also supported by dysregulated *c*REs are labeled.

To validate the regulatory function of the identified dysregulated *c*REs, we evaluated the genomic localization of DEGs with respect to *cis*-regulatory dysregulation in a cell type-specific manner. To this end, we integrated bulk RNA-seq dataset from 16 SN specimens (late-stage PD = 8, control = 8) with the snRNA-seq data to obtain a final set of 3,917 DEGs (down = 1,749, up = 2,168), which supplements the sparsity of snRNA-seq results (fig. S9 and table S5). Our colocalization analysis suggested that the DEGs were preferentially localized with dysregulated *c*REs in a 100kb genomic window compared to random expectations (Fig. 2B and fig. S10, A and B). In addition, GO analysis for DEGs and dysregulated *c*REs presented shared molecular pathways implicated in PD pathogenesis in a cell type-resolved manner (Fig. 2C). Taken together, the identification of genome-wide dysregulation in *cis*-regulatory landscape may explain the aberrant gene expression specific to PD.

### Identification of potential PD genes associated with dysregulated *c*REs and PD genetic risk variants

Given the strong link of dysregulated *cis*-regulatory regions with PD neuropathology, we aimed to systematically decipher the gene regulatory circuitry by implementing the “activity-by-contact” (ABC) model (*40*). It is an experimentally-proven model, in which the quantitative effect of a *c*RE over its target gene is determined by the activity of the *c*RE weighted by the interaction frequencies to the target gene (Fig. 3A). To this end, we performed *in situ* Hi-C experiment utilizing 11 SN specimens (late-stage PD = 5, control = 6), and sequenced a total of 5.16 billion mapped reads to obtain 1.61 billion valid *cis* read-pairs (744 million for PD SN and 870 million for control SN), generating unbiased, all-to-all 3D chromatin contact maps for PD and control SN (Fig. 3A and fig. S11A). Using Fit-Hi-C (*41*), we identified 1.42 million and 1.03 million long-range chromatin interactions in 5kb resolution within a 1Mbp window for PD and control SN, respectively (*Q* < 0.01; union = 1.87 million) (fig. S11B). The quality of the identified interactions was assessed by confirming the marked enrichment of these interactions at promoters (11.05%) and *c*REs (53.64%) (fig. S11, C and D). Additionally, the total chromatin contacts for a gene accounting all chromatin interactions anchored to its gene promoter significantly correlated with gene expression in all cell types (fig. S11E). Lastly, we experimentally validated promoter-to-*c*RE relationships identified based on significant interactions by mimicking a PD GWAS-SNP through CRISPR-Cas9-mediated genome disruption using SH-SY5Y neuroblastoma (Fig. 3B and fig. S12). The quantitative mRNA expression analysis on the genes (*TOMM7*, *KLHL7*, and *NUPL2*) linked to the PD GWAS-SNP harboring *c*RE, showed a significant reduction in gene expression after the induction of the genetic mutation (Fig. 3B).

**Fig. 3.**
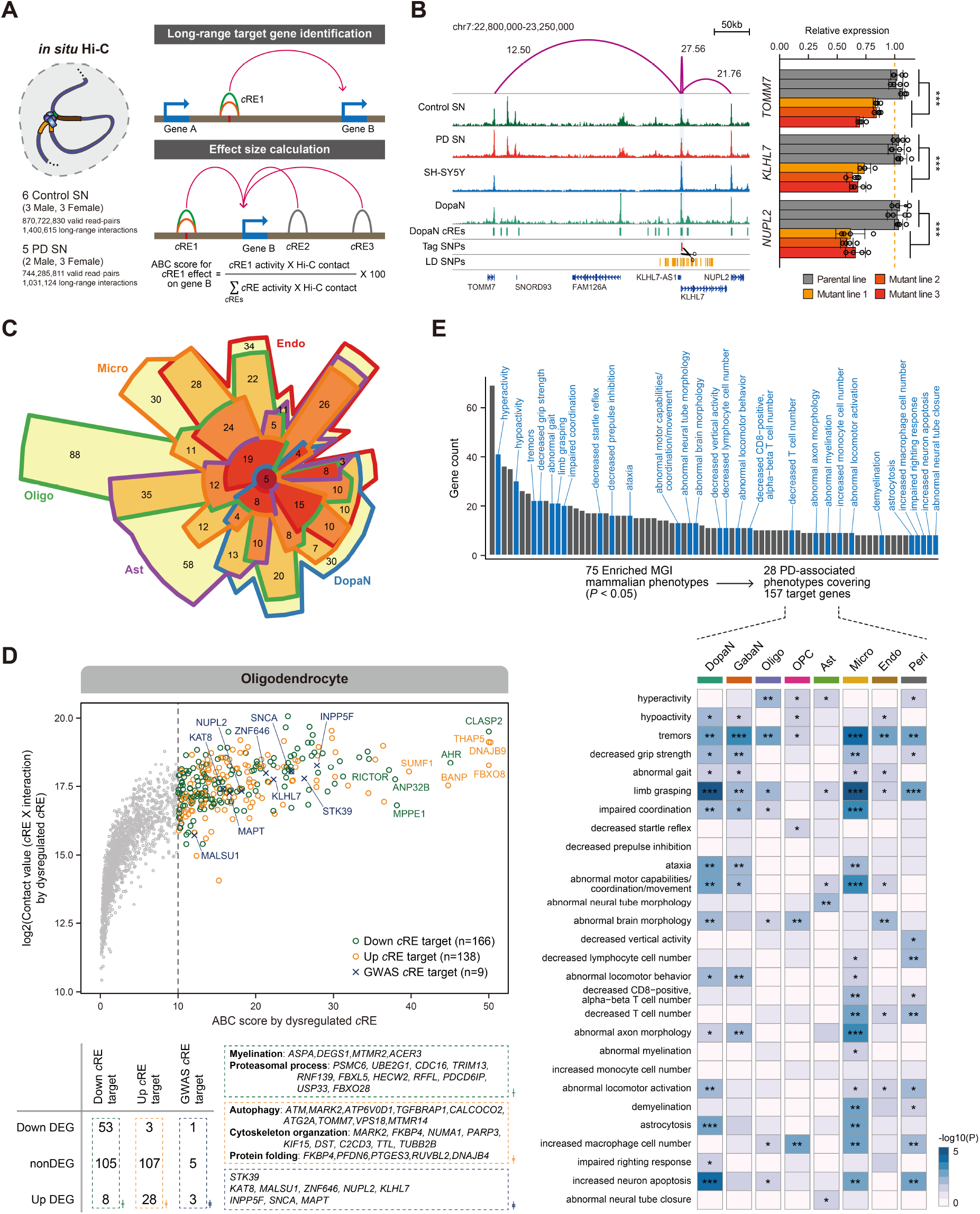
Target gene inference for non-coding regulatory sequences through 3D chromatin contacts. **(A)** Generation of high-resolution 3D chromatin contact map in the human SN. The illustrative approaches to identify long-range interaction target genes and to characterize the gene regulation circuitry by calculating the effect size of all *c*RE-to-gene promoter associations within 1Mb based on ABC scoring. **(B)** Left: H3K27ac ChIP-seq tracks for control SN, PD SN, and SH-SY5Y neuroblastoma, along with pseudobulk chromatin accessibility for dopaminergic neurons (DopaN). Additional tracks indicate positions of DopaN *c*REs and PD GWAS-SNPs. Significant long-range chromatin interactions shown in purple arcs with the corresponding ABC score. Guide RNA target sites for CRISPR-Cas9 mediated mutations were represented by scissors. The signals for ChIP-seq and cell type-resolved snATAC-seq were normalized by the total reads mapped in *c*REs. Right: Bar plots with dots indicating *TOMM7*, *KLHL7*, and *NUPL2* RNA levels in the parental SH-SY5Y cells and three independent mutant clones. Each clone having three biological replicates (Welch’s two-sample t-test, ***: p-value < 0.001). **(C)** A Chow-Ruskey plot for putative target genes of dysregulated *c*REs and PD GWAS-SNPs for dopaminergic neurons (DopaN), oligodendrocytes (Oligo), astrocytes (Ast), microglia (Micro), and endothelial cells (Endo). **(D)** Top: a scatter plot illustrating the putative target genes for oligodendrocyte with an ABC score (x-axis) threshold of 10, with the annotation of top 10 genes with the highest ABC score, along with target genes of GWAS-SNPs. Putative target genes were grouped into three categories, whether they are anchored by down- (green circles), up-regulated (orange circles) *c*REs, or GWAS-SNP harboring *c*REs (navy saltire). Bottom: putative target genes were categorized based on DEG status and the representative GO biological pathways. **(E)** Top: bar plots showing 75 enriched mammalian phenotypes (p-value < 0.05 and the number of associated genes ≥ 8) linked to putative targets of dysregulated *c*REs and PD GWAS-SNPs. 28 phenotypes associated with PD are highlighted in blue. Bottom: a heatmap illustrating the enrichment level of the 28 PD-associated mammalian phenotypes in individual cell types (*: p-value < 0.05, **: p-value < 0.01, ***: p-value < 0.001).

With the ABC model applied to our high-resolution chromatin contact map, we quantified all *c*RE-target gene relationships with respect to their contribution to target gene expression within 1Mb window. A total of 656 target genes of dysregulated *c*REs and PD GWAS-SNPs was identified in a cell type-specific manner (DopaN = 165, GabaN = 191, Oligo = 300, OPC = 231, Ast = 233, Micro = 223, Endo = 235, and Peri = 201; table S6) based on an ABC score threshold greater than 10 (equivalent to 10% contribution in overall chromatin contacts for a gene). These putative target genes were highly cell type-specific, with a considerable portion of the target genes (52.13%) assigned to only one or two cell types (Fig. 3C). The enriched biological pathways of up-regulated *c*REs indicated a recurrent representation of autophagy (DopaN with *SBF2* and *KLHL22*; Oligo with *MARK2*, *ATG2A*, and *TOMM7*; Ast with *WDR1* and *PPP1R9B*; Micro with *PTGES3* and *RUVBL2*) and protein folding (DopaN with *ABCA7*; Oligo with *PFDN6* and *DNAJB4*; Ast with *DNAJB5*) in multiple cell types (Fig. 3D and fig. S13). Conversely, cell type-specific biological pathways implicated in PD pathogenesis were enriched by the target genes of down-regulated *c*REs, including learning or memory (DopaN; *ATP8A1* and *ARL6IP5*), myelination (Oligo; *DEGS1*, *MTMR2*, and *ACER3*), brain morphogenesis (Ast; *BBS2* and *PAFAH1B1*), and protein lipidation (Micro; *MPPE1*, *ATG10*, and *ZDHHC20*) (Fig. 3D and fig. S13). Finally, a gene set enrichment analysis of putative PD genes based on MGI mammalian phenotype database showed a substantial association with PD-related phenotypes (28 of 75 phenotypes), involving 157 of 656 target genes (Fig. 3E, top). When the gene set analysis in these phenotypes was reiterated based on each cell type, we found varying cell types causing each of these phenotypes (Fig. 3E, bottom). Altogether, our findings strongly suggest that the identified putative target genes of dysregulated *c*REs and PD GWAS-SNPs have significant implications in PD neuropathology, and reaffirm that PD is a highly heterogeneous disorder with the involvement of diverse cell types and risk genes.

### Functional annotation of PD genetic risk variants in cell-type resolution

Based on the gene regulatory circuitry identified by 3D chromatin contact maps, a putative function of PD risk variants in the gene regulation program was annotated in individual cell types. To examine whether common genetic variants related to PD exert their function through modulating regulatory regions (*21, 42*), we collected and analyzed 5,912 genetic variants from four PD GWAS summary statistics (*P* < 5e-8) (*8, 9, 11, 13*). We found 61.69% of these genetic variants were associated with *c*REs by linkage disequilibrium (LD) (fig. S14). The enrichment of PD-related SNPs in cell type-resolved *c*REs was assessed by implementing linkage disequilibrium score (LDSC) regression analysis of heritability (*43, 44*). We incorporated 5 additional GWAS summary statistics of other neurological and psychiatric disorders (table S7) (*45–49*). Interestingly, oligodendrocyte *c*REs showed a strong association in three PD GWASs (Nalls et al. 2014, Chang et al. 2017, and Foo et al. 2017) (Fig. 4A). Microglia and endothelial *c*REs indicated an enrichment in two PD GWASs (Chang et al. 2017, and Foo et al. 2017), and dopaminergic neuron *c*REs were exclusively enriched in the PD GWAS conducted in East Asian cases (Foo et al. 2017). This finding shows that PD etiology involves more diverse cellular properties than AD, whose GWAS-SNPs are most heavily linked to microglia *c*REs (*23, 24, 34*). Further, the GWAS heritability analysis on dysregulated regulatory regions suggests that the PD risk variants were specifically enriched in down-regulated *c*REs (Fig. 4B), highlighting the association of genetic predisposition related to PD with down-regulated *cis*-regulatory landscape. For example, we found a down-regulated *c*RE in the vicinity of *SNCA*, which plays a critical role in PD pathogenesis (Fig. 4C). Finally, the depletion of genetic variants for other neurological and psychiatric disorders shows that PD has a distinct pathogenesis mechanism compared to these diseases, and that our dysregulated *c*REs represent aberrant regulatory features pertinent to PD susceptibility (Fig. 4B).

**Fig. 4.**
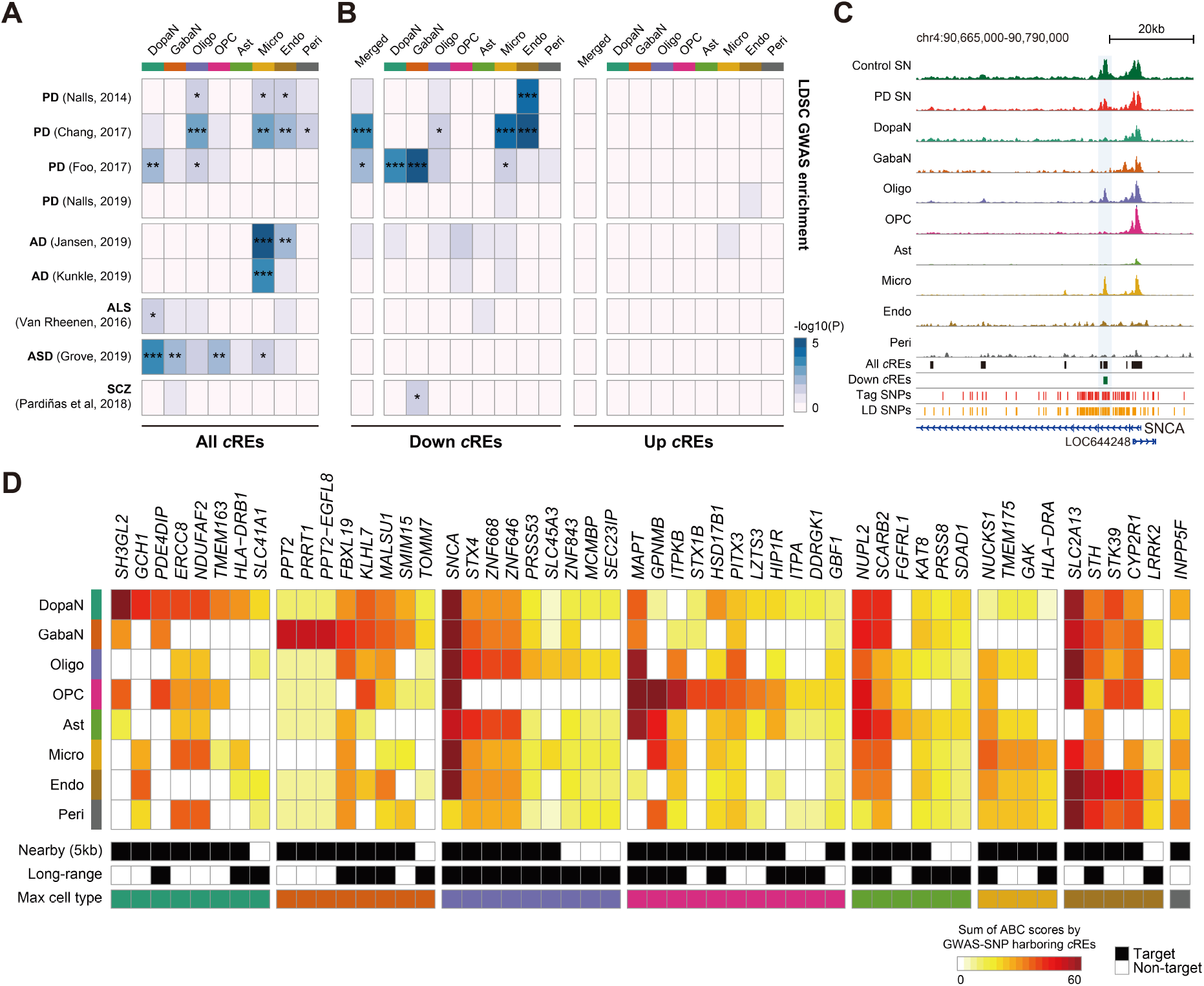
Characterization of PD GWAS-SNPs based on cell type-resolved epigenomic landscape. **(A-B)** Heatmaps illustrating the LDSC GWAS-SNP enrichment for neurological and psychiatric disorders in cell type-resolved *c*REs (A) and dysregulated *c*REs (B). AD, Alzheimer’s disease; ALS, amyotrophic lateral sclerosis; ASD, autism spectrum disorder; SCZ, schizophrenia. P-values were derived from the LDSC enrichment testing (*: p-value < 0.05, **: p-value < 0.01), ***: p-value < 0.001). **(C)** H3K27ac ChIP-seq tracks for PD and control SN, and cell type-resolved pseudobulk snATAC-seq signals in the *SNCA* locus. Additional tracks indicate the positions of all and down-regulated *c*REs, along with tag and LD-expanded PD GWAS-SNPs. The signals for ChIP-seq and cell type-resolved snATAC-seq were normalized by the total reads mapped in *c*REs. **(D)** A heatmap describing the putative target genes of GWAS-SNP harboring *c*REs identified based on significant chromatin interactions. The ABC scores are calculated iteratively based on the cell type-resolved epigenome to describe cell type-specific activation of key pathogenic genes by the PD GWAS-SNPs. 52 genes with the sum of ABC scores by the GWAS-SNP harboring *c*REs greater than 20 are shown.

Additionally, our target gene analysis based on 3D chromatin interactions and PD GWAS-SNPs unraveled important regulatory associations involving known PD genes, and annotated specific cell types in which these associations are active. For example, the functional association of PD GWAS-SNPs to *SH3HL2* and *GCH1* was highly specific to dopaminergic neurons, while *SNCA* was significant in most cell types (Fig. 4C). *MAPT* was dominantly associated in oligodendrocytes, oligodendrocyte precursor cells, and astrocytes. *GPNMB* was active in oligodendrocyte precursor cells, astrocytes, and microglia, while *SCARB2* was significantly associated in neurons and astrocytes (Fig. 4C). Our data substantially facilitate the functional interpretation of PD GWAS-SNPs, and link known risk factors of PD with the responsible cell types.

### Modular expression patterns of potential PD genes resolve complex molecular characteristics of PD

Since a number of potential PD genes has been identified by inferring the target genes of dysregulated *c*REs and PD GWAS-SNPs, we attempted to characterize the complex molecular pathways underlying the neuropathology based on the potential PD genes, and identify the responsible cell type associated with each pathway. To this end, we examined the dynamic modular gene expression patterns of the 656 potential PD genes utilizing the 16 bulk SN transcriptome from PD and control cases. The hierarchical clustering of gene expression correlation across these samples displayed a modular pattern with 9 distinct clusters (from C1 to C9) with notable biological processes involved in PD neuropathology (Fig. 5A and table S8). In addition, these biological pathways were represented by cell type-specific gene regulatory relationships. For example, genes involved in C1 (response to unfolded proteins, and reactive oxygen species) were specifically targeted by microglia *c*REs (Fig. 5B). Alternatively, the C3 genes represented by negative regulation of apoptosis were selectively enriched in oligodendrocytes and astrocytes, implicating their neuroprotective function toward the loss of nigral neurons. Surprisingly, C2 contained multiple cellular processes associated with PD pathogenesis, including endocytosis, lipid metabolism, iron homeostasis, and synaptic function (*50–53*), and harbored many target genes of PD GWAS-SNPs, highlighting the possibility that these cellular pathways are highly associated with PD pathogenesis (Fig. 5B). The genes in C2 were enriched with down-regulated *c*RE target genes from all cell types, which is consistent with our finding in the heritability analysis that PD-related genetic variations are associated with down-regulated *c*REs from diverse cell types. Furthermore, through our target gene analysis, we pin-pointed potential links to novel PD candidates while confirming the connections of known PD risk genes. For example, the endocytosis-related genes in C2 identified *CLASP2*, *PDCD6IP*, *MTMR2*, and *PICALM*, in addition to *SCARB2* and *INPP5F*, whose genetic variations are highly related to PD pathogenesis (Fig. 5, C to E) (*9, 54–56*).

**Fig. 5.**
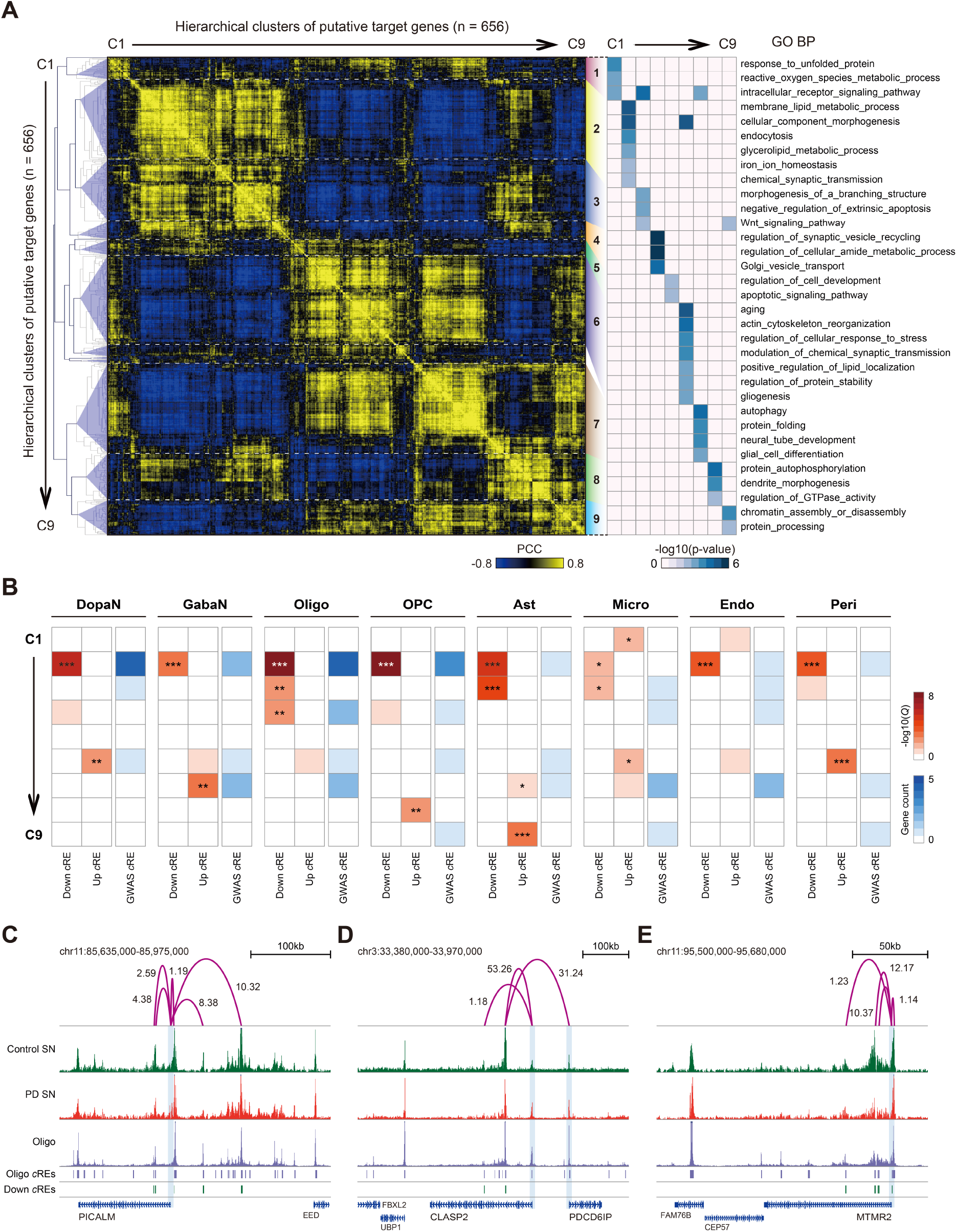
Analysis of modular gene expression patterns across putative PD genes. **(A)** Left: a hierarchical clustering of 656 putative PD genes based on similarities in gene expression between 16 bulk RNA-seq samples from control and PD SN. The color intensity indicates PCC of putative target gene expression between pair-wise samples. 9 distinct clusters were identified by a linkage distance (0.66) threshold in the dendrogram. Right: a heatmap describing the enrichment of GO biological pathways. Each entry indicates -log10(p-value) of GO biological processes in the corresponding cluster. **(B)** Heatmaps illustrating the enrichment of cell type-resolved target genes with respect to the 9 co-expression clusters. For each cell type, the putative target genes were categorized into three groups based on the type of *c*RE connected (down-, up-regulated *c*REs, or GWAS-SNP harboring *c*RE). The enrichment level was calculated based on one-sided exact binomial test with BH multiple testing correction (*: q-value < 0.05, **: q-value < 0.01, ***: q-value < 0.001). **(C-E)** H3K27ac ChIP-seq tracks for PD and control SN, and pseudobulk chromatin accessibility for oligodendrocytes (Oligo) for a genomic locus containing *PICALM* (C), *CLASP2*and *PDCD6IP* (D), and *MTMR2* (E). Additional tracks indicate positions of Oligo and down-regulated *c*REs. The signals for ChIP-seq and cell type-resolved snATAC-seq are normalized by the total reads mapped in *c*REs. Significant long-range chromatin interactions shown in purple arcs with the corresponding ABC score.

From C6 to C9, we found PD-associated biological features targeted by up-regulated *c*REs pertaining to a specific cellular identity (Fig. 5B). Importantly, we found that C6 represented fundamental pathogenic pathways (aging, cytoskeleton organization, stress response, modulation of synaptic activity, protein stability, and gliogenesis) related to dopaminergic neurons and microglia, while containing PD risk genes such as *MAPT*, *BAX*, and *TOMM7* (*57*). In C7, cellular processes such as glial cell differentiation and neural tube development were activated in astrocytes, in all likelihood, as a response mechanism to PD progression. Lastly, target genes in C8 indicate GTPase-mediated signaling and protein autophosphorylation are active in oligodendrocyte precursor cells as a result of autophagy accumulation in PD SN (*58, 59*).

Altogether, our analysis on the modular expression pattern demonstrates that the heterogeneous molecular features in the advanced PD cases are well explained by the target genes of dysregulated *c*REs and PD GWAS-SNPs. Although common genetic variations found in GWASs explain only a fraction of PD pathogenesis in individual PD cases, most of the PD-related cellular processes were represented by our target genes based on the integration of epigenetic features. The strong association between PD genetic variants and down-regulated *c*REs support the notion that cellular processes linked to PD are coordinated by the combined effects of genetic and epigenetic aberrations. Based on these findings, we propose that a genetic predisposition and epigenetic dysregulation are two indistinguishable modes of gene regulation that contribute to PD neuropathogenesis (fig. S15).

## Discussion

Major progresses have been made in the past decades furthering the understanding of molecular mechanisms and risk genes involved in PD pathogenesis. While evidences have revealed diverse PD-related cellular properties including protein quality control, autophagy-lysosome pathway, mitochondria homeostasis, lipid metabolism, synaptic toxicity, and neuroinflammation (*50–53*), our knowledge is still limited to fully explain the molecular causes of sporadic PD cases. A thorough characterization of the dynamic and interactive role of individual cell types during PD development is essential to broaden the scope of PD heritability. In this regard, the current study provides important insights into PD neuropathology by identifying the cell type-resolved dysregulation of *cis*-regulatory landscape, and by characterizing the perturbed molecular pathways based on the relevant cell types. Through this effort, we identified 656 novel PD candidate genes, which demonstrate the heterogenous molecular characteristics implicated in PD.

Our findings suggest that PD is, indeed, a highly heterogeneous disorder. The GWAS-SNP enrichment test of PD heritability showed that PD involves far more diverse cellular properties than AD, whose enrichment is limited predominantly in microglia. It is clear that the common variants identified by GWASs exhibit a regulatory function in a cell type-specific manner. However, most of previous genomics-based studies for complex traits did not sufficiently address this critical issue. We conducted cell type annotation of key genetic variants by overlaying them onto the cell type-resolved epigenome, and demonstrated that the PD variants are highly associated with genome regulatory elements and are likely conduct a regulatory role in cell type-specific manner, together with dysregulated *c*REs. In this regard, our results provide a unique perspective about the PD heritability that the previous genomics investigations were not able to address.

We employed the first systematic approach to quantitatively evaluate genome-wide *c*RE-target gene relationships using the ABC model. The implementation of ABC model to our high-resolution 3D contact maps defined the functional role of all *c*REs to the surrounding genes. The integrative analysis based on cell type-resolved epigenome revealed cell type functionality of key risk genetic variants and candidate genes, greatly advancing our view on the regulatory mechanisms involving PD neuropathology. The comprehensive evaluation of *c*RE-target gene connections also allowed the identification of potential PD candidates, by inferring high-confident target genes of dysregulated *c*REs and PD GWAS-SNPs based on a ABC score threshold. Our computational framework incorporating omics data integration is highly applicable in other complex human disorders.

This marks the first investigation, to our knowledge, to perform transcriptomic analysis cross-sectioning PD and control cases in the SN at single-nucleus level. We found that six of the known PD risk genes, including *SNCA* (Oligo, Micro), *MAPT* (Oligo, OPC), *UCHL1* (Oligo), *LRRK2* (Micro), *VPS13C* (Micro), and *CHCHD2* (Peri), are all up-regulated in their relevant cell types. It is interesting to notice that, while these risk genes are up-regulated, the PD variants are mostly associated with down-regulated *c*REs. This is consistent with the recent study suggesting a decoupling of promoter H3K27ac signals and gene expression in the PD brain (*22*). Nevertheless, our results suggest that the up-regulated DEGs and down-regulated *c*RE targets are involved in different cellular pathways. As shown in our modular expression profiling, down-regulated *c*RE and GWAS-SNP target genes are abundantly included in C2 (endocytosis, lipid metabolism, iron homeostasis, and synaptic function), while the up-regulated DEGs, such as *MAPT* and autophagy-related genes, are found in C6 (aging, cytoskeleton organization, stress response, modulation of synaptic activity, protein stability, and gliogenesis) and C7 (autophagy, protein folding, and glial differentiation). Our 3D epigenome analysis revealed specific *cis*-gene regulations that modulate these PD risk genes (*SNCA*, *MAPT*, *SCARB2*, *GCH1*, *BAG3*, and *INPP5F*), while identifying novel candidate genes that are associated with PD-implicated biological processes. The present work emphasizes the role of non-coding regulatory elements in understanding PD neuropathology, and provides additional insights into molecular mechanisms related to the perturbed epigenomic landscape.

The single-nucleus sequencing strategy is a highly advanced technique allowing the investigation of individual cell populations. However, due to the insufficient coverage obtained for each nucleus, the single-nucleus assays allow a differential analysis in only a small fraction of genes. The detection rate of DNA fragments is far less for snATAC-seq because of the limited copies of DNA to capture, in comparison to snRNA-seq. This inherent sparsity hinders robust identification of epigenomic dysregulation for a complex disease to its entirety. In this aspect, our work shows that the integration of bulk assays with a corresponding single-nucleus dataset may provide a solution. Our single-nucleus datasets offered a suitable reference for cellularity correction method for the bulk sequencing data, and the ratio of sequenced reads mapped to cell type markers effectively addressed the fraction of each cell type across the samples. Our cellularity-corrected bulk transcriptome confirmed a significant correlation with snRNA-seq, and allowed the identification of additional DEGs complementary to the single-nucleus dataset. We showed that the sequencing data generated from bulk tissues may still hold substantial value as a resource, when integrated with a matched single-nucleus data.

The difficulty in interpreting postmortem tissue data lies in the unresolved cause and effect signature by the pathology. Studies pin-pointing the epigenomic changes prior to the motor-phase of PD (Braak stage 1 and 2) or studies leveraging Braak stage-specific samples may further elucidate the molecular mechanism underlying the pathogenic progression of PD. Additionally, the chromatin conformation capture (3C) method conducted at a single-nucleus level will allow the cell type-specific investigation of chromatin interactome. The application of single-cell Hi-C technology (*60–62*) to clinical brain samples of neurodegenerative disorders may better portray the disease-related gene regulatory circuitry in a cell type-specific manner. Establishing the complete 3D epigenome may play an important part, especially, in effective personalized therapeutics, in light of the recent success in restoring clinical symptoms of PD by the implantation of patient-derived dopaminergic progenitor cells (*63*). Nevertheless, the present delineation of PD-specific aberration in *cis*-genome regulation, coupled with high-resolution chromatin interaction maps, significantly broadened the scope for disease-specific gene regulation mechanisms and expanded novel therapeutic candidate genes for PD. The present work conveying cell type-resolved non-coding regulatory elements lays the ground for further understanding of the gene regulatory network involved in the complex neuropathology.

## Materials and Methods

### Collection of human brain samples

Flash-frozen postmortem tissues from the human substantia nigra (SN) were acquired for PD (Braak stage = 3.51 ± 1.03) and control subjects (Braak stage = 0.66 ± 0.51) (table S1) from the Alzheimer Disease Research Center (ADRC) at University of California, San Diego. Samples from the left mid-frontal cortex were fixed in 4% PFA and the right were stored in liquid nitrogen for experiments. Formalin-fixed brains were sectioned at 40 μm, processed for pathohistological examination by Hematoxylin and Eosin (H&E), and a pathological scoring was assessed according to Braak stages (*64*). Institutional Review Board (IRB) approval was obtained from KAIST for the use of these brain tissues.

### Single-nucleus RNA and ATAC sequencing

About 15 mg of tissue was placed in an extra-thick tissue processing tube (Covaris, 520140) while kept in liquid nitrogen, and repeatedly hammered to produce frozen tissue powder. The frozen tissue was transferred to lysis buffer of 0.2% Triton X-100, protease inhibitor (Roche, 04-693-159-001), 1 mM DTT (Sigma-Aldrich, D9779), 0.2 U/µL RNAsin (Promega, N211B), and 2% BSA (Sigma-Aldrich, SRE0036) in PBS. Sample was pipetted 10 times, rotated for 5 min at 4°C, and centrifuged at 500g for 5 min. The pallet was resuspended in sorting buffer of 1 mM EDTA, 0.2 U/µL RNAsin, and 2% BSA in PBS, and passed through a 30 µm strainer (Sysmex, 04-0042-2316) to remove excessive debris. The nuclei were stained with DRAQ7 (1:100, Cell Signaling, 7406) for snRNA-seq, and with DAPI (10 ug/mL, Sigma, 32670) for snATAC-seq. Between 100,000 and 150,000 nuclei were sorted using a MoFlo Astrios EQ sorter (Beckman coulter, laser/filter) into a collection tube with 1 U/ µL RNAsin and 5% BSA in PBS. Sorted nuclei were centrifuged at 1000 g for 15 min at 4°C, and supernatant was removed. Nuclei were resuspended in resuspension buffer (0.2 U/µL RNAsin and 1% BSA in PBS) and counted using a cell counter (Cellometer Auto 2000, Nexcelom Biosicence) with AO/PI staining.

A subset of snRNA-seq libraries were generated by pooling the nuclei from three different donors with a matched pathological state, and demultiplexed based on individual genetic backgrounds using souporcell (*65*). We generated snRNA-seq libraries using the Chromium Single Cell 3’ Library & Gel Bead Kit v3 (10x Genomics). For snATAC-seq libraries, the isolated nuclei were subject to permeabilization in a lysis buffer with Tris-HCl (pH8.0), 10 mM NaCl, 3 mM MgCl_2_, 0.1% Tween-20, 0.1% Nonidet P40 Substitute (Sigma-Aldrich, 74385), 0.01% digitonin (Thermo Fisher Scientific, BN2006), and 1% BSA. After washing, the nuclei were resuspended in nuclei buffer, and snATAC-seq libraries were generated using the Chromium Single Cell ATAC Library & Gel Bead Kit v1 (10x Genomics). Quality control for DNA libraries was performed using Agilent Tape Station 4200 with high sensitivity D5000 kit. The libraries were sequenced in a paired-end mode using MGI DNBSEQ-G400 system.

### Single-nucleus RNA-seq data processing

In addition to the single-nucleus data that we generated from the SN specimens from ADRC, raw snRNA-seq data generated from healthy SN specimens (GSE140231) (*35*) was downloaded, and processed in parallel (table S1). To demultiplex the data for pooled libraries, the nuclei were clustered based on individual genetic variants using souporcell (*65*), and assigned to the corresponding donor by matching the genetic variants obtained from bulk RNA-seq data. The genetic variants from bulk RNA-seq data was obtained by using freebayes (-iXu -C 2 -q 20 -n 3 -E 1 -m 30 --min-coverage 20 --pooled-continuous --skip-coverage 100000). The feature-barcode matrix was generated using cellranger count (10x Genomics, v3.0.2) (*66*), aligning the sequenced reads to the human reference genome (hg19; 10x Cell Ranger reference GRCh37 v3.0.0). Then the count matrices were aggregated by cellranger aggr function across the samples with default parameters. The following analysis was performed using Seurat R package v4.0.5 (*67*). Nuclei with fewer than 200 or greater than 10,000 genes detected were filtered from the snRNA-seq dataset. Low-quality nuclei with mapped reads in the mitochondrial genes greater than 10% were removed. Doublets were identified using Scrublet (*68*), and excluded from analysis. The snRNA-seq data was integrated to correct for technical differences across individual samples. For this, the feature-barcode matrix was individually normalized by the total read count and log transformed, and top 5,000 variable genes were selected for each sample using Seurat’s FindVariableFeatures function. Data integration was conducted based on anchors identified by using FindIntegrationAnchors function. The aligned nuclei were scaled, and principal component analysis (PCA) was conducted on the scaled expression matrix. The top 50 principal components (PCs) were used to compute Shared Nearest Neighbors (SNNs), which were then used to cluster nuclei based on the Louvain algorithm (resolution = 1.5) in FindClusters function in Seurat R package. The top 50 PCs were used for the Uniform Manifold Approximation and Projection (UMAP) embedding.

Major cell types in the SN were assigned to each cluster based on known marker genes, including neurons (*SYT1*), oligodendrocyte (*MAG* and *MOBP*), oligodendrocyte precursor cells (*PDGFRA*), astrocyte (*AQP4* and *GFAP*), microglia (*CD74* and *RUNX1*), endothelial cells (*CLDN5*), and pericyte (*PDGFRB*). The nigral neurons were sub-clustered to identify dopaminergic and GABAergic neurons. We selected 1,000 highly variable genes by the *vst* method in Seurat R package v4.0.5 (*67*). Harmony (v1.0) (*69*) was used to correct the technical variations across the samples in the PCA dimensions. The first 10 dimensions were used to build SNN graph, which was clustered using the Louvain algorithm (resolution = 1.0). These dimensions were used to visualize neuronal nuclei in UMAP dimensions. Cell type markers for dopaminergic neurons (*TH* and *SLC6A3*) and GABAergic neurons (*GAD1* and *GAD2*) were used to assign sub-neuronal identity for individual sub-clusters. The nucleus-level expression signals were imputed using MAGIC (v2.0.3) (*70*) for UMAP visualization of cell type markers in snRNA-seq clusters. To build a reference for cell type-dependent transcriptome in the SN, we generated a count matrix from the feature-barcode matrix using the cellular identity annotated for all nuclei. The count matrix was then quantile-normalized, and the ratio of normalized reads based on individual cell types was computed to represent the gene expression ratio among different cell types. The cell types whose gene expression ratio is greater than 10% were annotated as the active cell type for the corresponding gene.

### Single-nucleus ATAC-seq data processing

In addition to the single-nucleus data we generated from the SN specimens from ADRC, raw snATAC-seq data generated from postmortem SN specimens (GSE147672) (*23*) was downloaded, and processed in parallel (table S1). The feature-barcode matrix was generated using the cellranger-atac count (10x Genomics, v1.1.0) (*71*), aligning the sequenced reads were aligned to the human reference genome (hg19). Then the count matrices were aggregated by cellranger-atac aggr function with default parameters. The following analysis was performed using Signac R package v1.4.0 (*72*). Nucleosome signal was computed with Signac’s NucleosomeSignal function. Low-quality nuclei were removed from snATAC-seq dataset based on the following criteria: fewer than 2,000 or greater than 30,000 fragments mapped to peak regions, less than 15% of reads in peak regions, nucleosome signal greater than 10, and transcription start site (TSS) enrichment less than 2. Doublets were identified using Scrublet (*68*), and excluded from analysis. We selected top 50% most common features as the variable features, and performed LSI dimensionality reduction by implementing term frequency-inverse document frequency (TF-IDF) transformation followed by singular value decomposition (SVD). Then, reciprocal LSI projection was conducted to identify integration anchors for each sample, and the snATAC-seq data was integrated across the samples by using low-dimensional cell embeddings with Signac’s IntegrateEmbeddings function. The identical method used in snRNA-seq for graph-based clustering and non-linear dimension reduction by UMAP was applied to the snATAC-seq dataset. Gene activity scores were computed for protein coding genes by summing snATAC-seq reads mapped in the gene body and the promoter (5 kb upstream to TSS) by using GeneActivity function in Signac R package.

Major cell types in the SN were assigned to each snATAC-seq cluster based on the identical set of known marker genes used in snRNA-seq processing. The neuronal population was sub-clustered to identify dopaminergic neurons and GABAergic neurons. First, we selected features with fragments detected in more than 10 nuclei as the variable features. Then, harmony (v1.0) (*69*) was used to correct the technical variations across the samples in the LSI dimensions. The first 5 dimensions were used to build SNN graph, which was clustered using the Louvain algorithm (resolution = 1.0). These dimensions were used to visualize neuronal nuclei in UMAP dimensions. Cell type markers for dopaminergic neurons (*TH* and *SLC6A3*) and GABAergic neurons (*GAD1* and *GAD2*) were used to assign sub-neuronal identity for individual sub-clusters. The nucleus-level gene activity scores were imputed using MAGIC (v2.0.3) (*70*) for UMAP visualization of cell type markers in snATAC-seq clusters.

We generated cell type-resolved BAM files for each sample, and merged the BAM files according to the cell type. Peak calling by MACS2 (*73*) identified 257,458 peaks (*P* < 0.05). Through a manual inspection of pseudobulk signals in the epigenome browser, we collected top 50% peaks (n = 128,724) based on the abundance of snATAC-seq reads, and defined them as *cis*-regulatory elements (*c*REs). To build a reference for cell type-dependent epigenome in the SN, we generated a count matrix by conducting bedtools (v2.29.1) (*74*) coverage function for the cell type-resolved BAM files. The count matrix was then quantile-normalized, and the ratio of normalized reads based on individual cell types was computed to represent the *c*RE activity among different cell types. The cell types whose *c*RE activity ratio is greater than 10% were annotated as the active cell type for the corresponding *c*RE.

### Identification of cell type-dependent differential features

We used FindAllMarkers function in Seurat R package to identify cell type markers with the model-based analysis of single-cell transcriptomics (MAST) algorithm (*75*) based on the two data modalities, leveraging the gene expression levels (snRNA-seq) and gene activity scores (snATAC-seq). It computes differential genes by iteratively contrasting one cell type to all, and the genes that satisfy Benjamini & Hochberg (BH)-adjusted *P* < 0.05, log2 fold change > 0.25, and expression detected in at least 10% of nuclei were defined as cell type markers. Because a subset of cell type markers was present in multiple cell types, we further identified unique cell type markers, in which the gene is identified as cell type marker in one cell type. In addition, we computed differentially expressed genes (DEGs), which was conducted iteratively for individual cell types comparing nuclei obtained from PD and control subjects using the snRNA-seq dataset. Seurat’s FindMarkers function was used with the MAST algorithm to identify cell type-specific DEGs that satisfy the following criteria: 1) BH-adjusted *P* < 0.05, 2) log2 fold change > 0.2, and 3) expression detected at least 10% of nuclei for both PD and control groups.

### Bulk RNA sequencing and data processing

About 40 mg of tissue was placed in an extra-thick tissue processing tube (Covaris, 520140) while kept in liquid nitrogen, and repeatedly hammered to produce frozen tissue powder. Total RNA was extracted using NucleoSpin RNA XS (Macherey-Nagel, 740902). RNA-seq libraries were prepared using TruSeq stranded mRNA library prep kit (Illumina, 20020594). External RNA controls consortium (ERCC) RNA spike-in mixes (Thermo Fisher Scientific, 4456740) were included for quality assurance. RNA-seq libraries were sequenced in a paired-end mode using Illumina HiSeq 4000 platform.

Paired-end reads were aligned to the reference genome (hg19 with ERCC) using STAR software v2.7.5 (*76*) with default parameters. The raw read counts were quantified with RSEM (*77*) based on a gene list obtained from GENCODE v38 by selecting protein coding genes and long non-coding RNAs (lncRNAs) with confidence level 1 and 2 (n=21,151). The count values were merged into a count matrix, and quantile normalized using preprocessCore R package across the samples. The cellular heterogeneity in individual samples was assessed with unique marker genes, and the overall gene expression pattern was adjusted iteratively based on relative gene expression ratios across the cell types. Technical variations from experimental and sequencing batches were corrected using ComBat (*78*) function in sva R package. We used the quasi-likelihood F-test in EdgeR (*39*) to identify differentially expressed genes (DEGs) with a multi-test corrected (BH method) significance threshold (adjusted *P* < 0.05). We annotated active cell types for each DEG based on the cell type reference obtained from single-nucleus RNA-seq data. After evaluating the correlation between cell type-resolved DEGs obtained from bulk RNA-seq and snRNA-seq, we combined these two DEG sets in a cell type-dependent manner, independently for down- and up-regulated DEGs.

### Chromatin immunoprecipitation sequencing

We conducted chromatin immunoprecipitation sequencing (ChIP-seq) to profile the genome-wide landscape of histone 3 lysine 27 acetylation (H3K27ac). About 40 mg of tissue was placed in an extra-thick tissue processing tube (Covaris, 520140) while kept in liquid nitrogen, and repeatedly hammered into frozen tissue powder. The tissue sample was crosslinked in a crosslinking buffer of 100mM NaCl, 0.1mM EDTA, 5mM HEPES (pH 8.0), and 1% formaldehyde for 10 min at room temperature. The crosslinking was quenched with 125 mM glycine for 5 min on a rotation, and washed twice with ice-cold PBS. The samples were passed through a 30 µm strainer (Sysmex, 04-0042-2316) to remove excessive debris, and suspended in SDS lysis buffer of 1% SDS, 50 mM Tris-HCl (pH8.0), 10 mM EDTA and protease inhibitor (Roche, 04-693-159-001). Chromatin fragmentation was performed by sonication (Covaris, S220) in the volume of 100ul to obtain mono-, di-, and tri-nucleosome size chromatin. After centrifugation at 12,000g for 15 min at 4°C, the sonicated chromatin in supernatant was diluted 10 times with dilution buffer to achieve final concentration of 0.1% Triton X-100, 0.1% SDS, mM NaCl, 15 mM Tris-HCl (pH 8.0), 1 mM EDTA, and protease inhibitor for chromatin immunoprecipitation (ChIP). The sonicated chromatin in supernatant was incubated with protein Dynabead (Thermo Fisher Scientific, 10001D) coated with anti-H3K27ac antibody (Active Motif, 39133) for 4 hours at 4°C with rotation, while a fraction of the input chromatin was stored to be used as an input control. The chromatin-antibody-bead complex was subjected to serial washing with varying salt concentrations optimized for the antibody used. The immunoprecipitated complex was treated with RNaseA (QIAGEN, 19101), and reverse-crosslinked overnight at 68°C. The immunoprecipitated DNA was recovered using AMPure XP beads (Beckman Coulter, A63881), and ChIP-seq libraries were prepared using NEBNext Ultra II DNA library Prep Kit (NEB, E7645) following the manufacturer’s instruction. The ChIP-seq libraries were sequenced in paired-end mode using Illumina HiSeq 4000 platform.

### Quantification of *cis*-regulatory element activity

The sequenced DNA reads from ChIP-seq libraries were mapped to human reference genome (hg19) using Burrows-Wheeler aligner (BWA)-mem (*79*) (ver. 0.7.17, ‘-M’ option). Reads with a low alignment quality (MAPQ < 10) were removed, and PCR duplicates were discarded using Picard (v2.6.0). We computed the number of ChIP-seq reads aligned in the 128,724 *c*REs identified based on snATAC-seq data using bedtools (v2.29.1) (*74*) coverage function. The read counts were merged into a count matrix, and quantile normalized using preprocessCore R package across the samples. The cellular heterogeneity in individual samples was assessed with unique marker *c*REs, and the global *c*RE activities were adjusted iteratively based on relative *c*RE activity ratios across the cell types. Technical variations from experimental and sequencing batches were corrected using ComBat (*78*) function in sva R package. We used the quasi-likelihood F-test in EdgeR (*39*) to identify dysregulated *c*REs with a multi-test corrected (BH method) significance threshold (adjusted *P* < 0.05). We annotated active cell types for each dysregulated *c*RE based on the cell type reference obtained from single-nucleus ATAC-seq data.

### Assessment and adjustment of cellular heterogeneity

To investigate the cellular heterogeneity in the bulk RNA-seq data generated from the SN tissues, we used 100 unique cell type marker genes for each cell type based on the cell type-resolved transcriptome identified by snRNA-seq data. For individual samples, we computed the sum of reads mapped to the marker genes from each cell type, and the ratio of these summed reads across cell types to evaluate the relative composition of cell populations within the bulk data. The identified composition of reads in the unique marker genes considerably matched the cellular compositions obtained from single-nucleus sequencing datasets. For each cell type, we calculated the mean of relative compositions computed from the bulk samples, and then a relative cellular fraction (RCF) was obtained for a given cell type in a sample by dividing its relative composition to the mean. Then, cellularity-adjusted value (CAV) for a gene *i*, was computed based on the RCF of a sample and the expression ratio (ER) among 8 cell types present in the SN as the following:

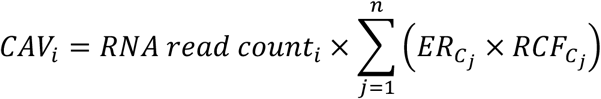

Where {*C_j_*} = {*DopaN*, *GobaN*, *Oligo*, *OPC*, *Ast*, *Micro*, *Endo*, and *Peri*}. This procedure was repeated three times until the variation of cellular compositions was minimal across the samples.

To examine the cellular heterogeneity in the bulk H3K27ac ChIP-seq data, we used top 200 unique cell type marker *c*REs for each cell type based on the cell type-resolved chromatin accessibility identified by snATAC-seq data. Identical to the method applied to bulk RNA-seq dataset, we computed the ratio of summed reads mapped to cell type marker *c*REs for individual samples, and the mean of relative compositions was calculated for each cell type. Then, the RCF was calculated for each cell type in a sample, and CAV for a *c*RE *i* was computed based on the RCF of the sample and the *c*RE activity ratio (CR) among 8 cell types present in the SN as the following:

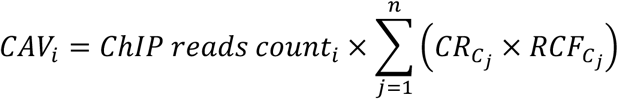

Where {*C_j_*} = {*DopaN*, *GobaN*, *Oligo*, *OPC*, *Ast*, *Micro*, *Endo*, and *Peri*}. This procedure was repeated three times until the variation of cellular compositions was minimal across the samples.

### Co-localization of DEGs in proximity of dysregulated *c*REs

To examine the regulatory effect of dysregulated *c*REs to the surrounding genes, we conducted an enrichment test to measure the number of DEGs harbored by the dysregulated *c*REs in a given genomic window (100kb), compared to random expectations. We created two sets of random groups, simulating both dysregulated *c*REs and DEGs. First, simulated *c*REs were generated by creating a set of genome coordinates that match the dysregulated *c*REs in number, size, and chromosome. Simulated DEGs were created by random gene sampling from the total gene set. The enrichment was measured based on iterative trials (n = 10,000) considering the degree of DEGs colocalized in 100kb window. The statistical significance was calculated in the form of empirical testing. The test was performed independently with respect to cell types and the type of *c*RE dysregulation.

### PD GWAS-SNP imputation based on LD structure

PD-related GWAS-SNPs were collected from four GWAS summary statistics (*8, 9, 11, 13*), and tag GWAS-SNPs with the significance threshold (*P* < 5e-8) were selected for down-stream analysis. We expanded the GWAS-SNPs using linkage disequilibrium (LD) structure. LD scores were calculated using PLINK for five different populations, including African (AFR), American (AMR), East Asian (EAS), European (EUR), and South Asian (SAS), from 1000 genome phase 3 data. For each tag SNP, we identified associated SNPs that share a tight LD score (r^2^ > 0.8) in at least three ethnic groups.

### LD score regression of disease heritability

To determine whether cell type-resolved *c*REs and dysregulated *c*REs are enriched with heritability of specific neurological and psychiatric disorders, we applied linkage disequilibrium score (LDSC) regression analysis (*43, 44*). We used four GWAS summary statistics for PD (*8, 9, 11, 13*). We obtained five additional GWAS summary statistics for Alzheimer’s disease (AD) (*45, 46*), amyotrophic lateral sclerosis (ALS) (*47*), autism spectrum disorder (ASD) (*48*), and schizophrenia (SCZ) (*49*). The cell type-resolved *c*REs and dysregulated *c*REs were tested for enrichment of heritability while controlling for the full baseline model.

### *In situ* Hi-C library preparation and data processing

*In situ* Hi-C experiments were performed on the SN tissues from PD and control cases (table S1). About 50 mg tissue was placed in an extra-thick tissue processing tube (Covaris, 520140), and repeatedly hammered while frozen in liquid nitrogen to produce tissue powder. The pulverized tissue was crosslinked with 1% formaldehyde. *In situ* Hi-C was conducted based on previously reported protocol with minor modifications (*33*). Crosslinked cells were lysed with 10nM Tris-HCl (pH8.0), 10mM NaCl, and 0.2% IGEPAL CA630 (Sigma-Aldrich, 18896), and digested with 100U MboI (NEB, R0147). Digested fragments were labelled with biotin-14-dCTP (Invitrogen, 19518018) and proximally ligated with T4 DNA Ligase (NEB, M0202), followed by reverse-crosslinking with 2 ug/uL proteinase K (NEB, P8102), 1% SDS, and 500 mM NaCl overnight at 68°C. The DNA fragments were purified with AMPure XP beads (Beckman Coulter, A63880), and subjected to sonication (Covaris, S220). The ligated DNA fragments were pulled down with Dynabeads MyOne streptavidin T1 beads (Invitrogen, 65602) with thorough washing. Hi-C libraries were prepared manually by performing DNA end repair, removal of un-ligated ends, adenosine addition at 3’ end (NEB, M0212), ligation of Illumina indexed adapters (NEB, M2200), and PCR amplification (Thermo Fisher Scientific, F549). The number of cycles for PCR amplification was determined based on KAPA library quantification kit (KAPA, KK4854). The Hi-C libraries were then subjected to deep sequencing in paired-end mode using Illumina HiSeq 4000 and X platforms.

The sequencing output from Hi-C libraries were mapped to the reference genome (hg19) using BWA-mem (*79*) (‘–M’ option). In-house scripts were used to remove low quality reads (MAPQ < 10), the reads that span ligation sites, chimeric reads, and self-interacting reads (two fragments located within 5kb). The chimeric reads were removed since they are bi-products of ligation chemistry in Hi-C library construction, and cannot be properly processed by paired-end BWA-mem command. The read-pairs were merged together as paired-end aligned BAM files, and PCR duplicates were removed with Picard (v2.6.0).

### Significant chromatin interaction calling

Statistically significant, long-range chromatin interactions were identified at 5kb resolution using Fit-Hi-C, as previously described (*41*). We created merged Hi-C BAM files with respect to control and PD status, and converted them into an input format for Fit-Hi-C in each chromosome, and used the default Fit-Hi-C code to calculate the interaction significance between two genomic coordinates in 1Mbp distance. A significance threshold (q-value < 0.01) was used to define significant chromatin interactions. We defined the union of chromatin interactions obtained from PD and control SN as a general interaction set, and classified promoter- and *c*RE-associated chromatin interactions by determining whether either bin of a chromatin interaction contained a transcription start site (TSS) or a *c*RE. We labeled them as ‘none’ if the bin in the chromatin interaction contained no regulatory element. Following this scheme, we defined the distal chromatin contacts into six categories: promoter-*c*RE (P-*c*RE), promoter-promoter (P-P), *c*RE-*c*RE, promoter-none (P-none), *c*RE-none, and none-none chromatin interactions.

### Calculation of activity by contact score

We applied a conceptually identical framework described in the activity by contact (ABC) model (*40*), in which the quantitative effect of a *c*RE to a target gene depends on the frequency with which it contacts its promoter multiplied by the activity of the *c*RE. Briefly, ABC score for effect of a *c*RE *i* on gene *j* was measured as the following:

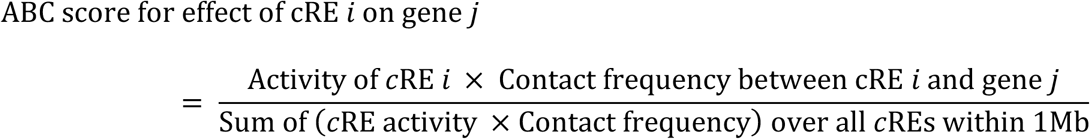

The ABC scores for all *c*RE to gene relationships within 1Mb window were computed for individual cell types. The *c*RE activity was defined by the normalized snATAC-seq reads in the given cell type, and the contact frequency was represented by Hi-C contact frequency between the two bins containing the *c*RE and TSS of the target gene in 5kb resolution. The position of a *c*RE was determined by the 5kb genomic bin, in which the center of the cRE was located. When two *c*REs were present in one 5kb bin, the sum of these *c*REs were used. Contact value was defined by *c*RE activity multiplied by contact frequency.

### Putative target gene identification for dysregulated *c*REs and PD GWAS-SNPs

Putative target genes of dysregulated *c*REs were identified iteratively for each cell type based on following criteria: (1) dysregulated *c*RE and its target gene are connected by a significant interaction, (2) dysregulated *c*RE and its target gene are annotated as active cell type in the cell type-resolved transcriptomic and epigenomic reference, (3) ABC score of dysregulated *c*RE to target gene relationship is greater than 10, and (4) the Pearson’s correlation coefficient (PCC) between *c*RE activity and target gene expression across the samples based on bulk sequencing data is greater than 0.3. Putative target genes of PD GWAS-SNPs were identified for individual cell types based on following data: (1) SNP harboring *c*RE and its gene are annotated as active cell type in the cell type-resolved transcriptomic and epigenomic reference, (3) ABC score of SNP harboring *c*RE to target gene relationship is greater than 10.

### Gene ontology and mammalian phenotype analyses

EnrichR (*80*) was used to identify biological processes for cell type-specific DEGs based on GO Biological Processes 2018 database. Similarly, GREAT (*81*) was used to determine biological processes that are associated with dysregulated *c*REs. We used enrichR to identify mammalian phenotype over-represented in putative target genes of dysregulated cREs and PD GWAS-SNPs based on 2021 mammalian phenotype level 4 database from mouse genome informatics (MGI) (*82*). Phenotypes with no biological relevance to human diseases, including ‘lethality’, ‘death’, and ‘no abnormal phenotype’ were excluded. Phenotypes with number of associated genes at least 8 and significance threshold (*P* < 0.05) were selected.

### CRISPR-Cas9 genome editing

We validated gene-to-regulatory sequence relationships identified by our Hi-C chromatin interactions. CRISPR-Cas9-mediated genome editing was conducted using ribonucleoprotein (RNP) delivery method in the SH-SY5Y cell-line. Three crRNAs, along with tracrRNA, were synthesized in vitro (IDT). To create a RNP complex, a guide and tracrRNA were annealed, mixed with Cas9 nuclease (Enzynomics, M058HL), and incubated for 15 min at room temperature. The RNP complex was transfected into cells by electroporation with Neon transfection 10 µl kit (Thermo Fisher Scientific, MPK1096). To measure the efficiency of genome editing in each guide RNA, we performed targeted deep sequencing. For this, the genomic DNA was extracted from the transfected cells and the target sites were amplified by PCR subsequently. Indices and sequencing adaptors were attached by additional PCR. High-throughput sequencing was performed using Illumina Mini-seq (San Diego, CA, USA). The mutation frequencies and patterns were analyzed using the Cas-Analyzer program implemented in CRISPR RGEN Tools (https://www.rgenome.net/). The cells were separated into single clones by serial dilutions on 96-well plates. After sufficient growth of each clone, the genotype was confirmed by conducting a Sanger sequencing of the target region from both directions. We selected mutant clones with the largest mutation size from each of the three guide RNAs from Sanger-sequencing results, and purified total RNA for reverse transcriptase quantitative PCR (RT-qPCR) to measure the relative mRNA expression levels of putative target genes.

crisprRNA #1: TCTTGTGTGAAGAAACCCGTTGG
crisprRNA #2: GCCCAAACCGAAGCCCCCAAAGG
crisprRNA #3: AGCAACTCTCCTCCCTTTGGGGG
Genotyping F: TCGTCTGCCGAGGATGTA
Genotyping R: AATTTCACGAATGCACCACAC
RT-qPCR primer (*GAPDH* F): CCACTCCTCCACCTTTGACG
RT-qPCR primer (*GAPDH* R): TTCGTTGTCATACCAGGAAATGAG
RT-qPCR primer (*TOMM7* F): CGGAATGCCTGAACCAACT
RT-qPCR primer (*TOMM7* R): GCCTTGTGCCATCCAACTA
RT-qPCR primer (*KLHL7* F): CAGCAAGAAGAAGACCGAGAAG
RT-qPCR primer (*KLHL7* R): GCAAGAACAACACGATGAGCAG
RT-qPCR primer (*NUPL2* F): AAGTTTGGGAGTCGTCGGGA
RT-qPCR primer (*NUPL2* R): CTTTTACGTCAGAGAGCAGAGC

### Analysis of modular gene expression patterns

For the 656 putative target genes were obtained, a correlation matrix where each entry indicates a similarity score between two putative target genes by computing the PCC based on 16 bulk RNA-seq samples. The correlation matrix was subjected to a hierarchical clustering (Pearson correlation metric with average linkage), which presented 9 distinct gene clusters at a dendrogram height threshold of 0.65. Enriched biological processes of protein-coding genes in clusters from C1 to C9 were determined using metascape (v3.5) (*83*). Enrichment of target genes based on the cell type and the type of dysregulated *c*REs in each cluster was evaluated using the one-sided exact binomial test. The corresponding significance values were multiple testing corrected for the number of cell type annotations.

## Acknowledgments

We thank the members of Jung laboratory and Dr. Andrew Singleton for support and critical suggestions throughout the course of this work.

## Author contributions

A.J.L., C.K., E.M., and I.J. conceived the study. A.J.L., S.P., and J.E. performed sequencing library preparation. A.J.L. and K.J. performed molecular experiments. A.J.L. conducted bioinformatics analysis. C.K. and E.M. provided human brain samples. C.K. and R.A.R. performed histopathologic quality control for clinical specimens. C.K., E.M., S.J.L., S.J.C., and J.C. contributed to result interpretation. A.J.L., and C.K. prepared the manuscript with assistance from E.M. and I.J. All authors read and commented on the manuscript.

## References and Notes

1. J. Jankovic, Parkinson’s disease: clinical features and diagnosis. J Neurol Neurosurg Psychiatry 79, 368–376 (2008).

2. A. Berardelli, J. C. Rothwell, P. D. Thompson, M. Hallett, Pathophysiology of bradykinesia in Parkinson’s disease. Brain 124, 2131–2146 (2001).

3. N. Caballol, M. J. Marti, E. Tolosa, Cognitive dysfunction and dementia in Parkinson disease. Mov Disord 22 Suppl 17, S358–366 (2007).

4. J. Park, S. B. Lee, S. Lee, Y. Kim, S. Song, S. Kim, E. Bae, J. Kim, M. Shong, J. M. Kim, J. Chung, Mitochondrial dysfunction in Drosophila PINK1 mutants is complemented by parkin. Nature 441, 1157–1161 (2006).

5. A. B. Singleton, M. Farrer, J. Johnson, A. Singleton, S. Hague, J. Kachergus, M. Hulihan, T. Peuralinna, A. Dutra, R. Nussbaum, S. Lincoln, A. Crawley, M. Hanson, D. Maraganore, C. Adler, M. R. Cookson, M. Muenter, M. Baptista, D. Miller, J. Blancato, J. Hardy, K. Gwinn-Hardy, alpha-Synuclein locus triplication causes Parkinson’s disease. Science 302, 841 (2003).

6. T. Kitada, S. Asakawa, N. Hattori, H. Matsumine, Y. Yamamura, S. Minoshima, M. Yokochi, Y. Mizuno, N. Shimizu, Mutations in the parkin gene cause autosomal recessive juvenile parkinsonism. Nature 392, 605–608 (1998).

7. V. Bonifati, P. Rizzu, M. J. van Baren, O. Schaap, G. J. Breedveld, E. Krieger, M. C. Dekker, F. Squitieri, P. Ibanez, M. Joosse, J. W. van Dongen, N. Vanacore, J. C. van Swieten, A. Brice, G. Meco, C. M. van Duijn, B. A. Oostra, P. Heutink, Mutations in the DJ-1 gene associated with autosomal recessive early-onset parkinsonism. Science 299, 256–259 (2003).

8. D. Chang, M. A. Nalls, I. B. Hallgrimsdottir, J. Hunkapiller, M. van der Brug, F. Cai, C. International Parkinson’s Disease Genomics, T. andMe Research, G. A. Kerchner, G. Ayalon, B. Bingol, M. Sheng, D. Hinds, T. W. Behrens, A. B. Singleton, T. R. Bhangale, R. R. Graham, A meta-analysis of genome-wide association studies identifies 17 new Parkinson’s disease risk loci. Nat Genet 49, 1511–1516 (2017).

9. M. A. Nalls, N. Pankratz, C. M. Lill, C. B. Do, D. G. Hernandez, M. Saad, A. L. DeStefano, E. Kara, J. Bras, M. Sharma, C. Schulte, M. F. Keller, S. Arepalli, C. Letson, C. Edsall, H. Stefansson, X. Liu, H. Pliner, J. H. Lee, R. Cheng, C. International Parkinson’s Disease Genomics, G. I. Parkinson’s Study Group Parkinson’s Research: The Organized, andMe, GenePd, C. NeuroGenetics Research, G. Hussman Institute of Human, I. Ashkenazi Jewish Dataset, H. Cohorts for, E. Aging Research in Genetic, C. North American Brain Expression, C. United Kingdom Brain Expression, C. Greek Parkinson’s Disease, G. Alzheimer Genetic Analysis, M. A. Ikram, J. P. Ioannidis, G. M. Hadjigeorgiou, J. C. Bis, M. Martinez, J. S. Perlmutter, A. Goate, K. Marder, B. Fiske, M. Sutherland, G. Xiromerisiou, R. H. Myers, L. N. Clark, K. Stefansson, J. A. Hardy, P. Heutink, H. Chen, N. W. Wood, H. Houlden, H. Payami, A. Brice, W. K. Scott, T. Gasser, L. Bertram, N. Eriksson, T. Foroud, A. B. Singleton, Large-scale meta-analysis of genome-wide association data identifies six new risk loci for Parkinson’s disease. Nat Genet 46, 989–993 (2014).

10. C. International Parkinson Disease Genomics, M. A. Nalls, V. Plagnol, D. G. Hernandez, M. Sharma, U. M. Sheerin, M. Saad, J. Simon-Sanchez, C. Schulte, S. Lesage, S. Sveinbjornsdottir, K. Stefansson, M. Martinez, J. Hardy, P. Heutink, A. Brice, T. Gasser, A. B. Singleton, N. W. Wood, Imputation of sequence variants for identification of genetic risks for Parkinson’s disease: a meta-analysis of genome-wide association studies. Lancet 377, 641–649 (2011).

11. J. N. Foo, L. C. Tan, I. D. Irwan, W. L. Au, H. Q. Low, K. M. Prakash, A. Ahmad-Annuar, J. Bei, A. Y. Chan, C. M. Chen, Y. C. Chen, S. J. Chung, H. Deng, S. Y. Lim, V. Mok, H. Pang, Z. Pei, R. Peng, H. F. Shang, K. Song, A. H. Tan, Y. R. Wu, T. Aung, C. Y. Cheng, F. T. Chew, S. H. Chew, S. A. Chong, R. P. Ebstein, J. Lee, S. M. Saw, A. Seow, M. Subramaniam, E. S. Tai, E. N. Vithana, T. Y. Wong, K. K. Heng, W. Y. Meah, C. C. Khor, H. Liu, F. Zhang, J. Liu, E. K. Tan, Genome-wide association study of Parkinson’s disease in East Asians. Hum Mol Genet 26, 226–232 (2017).

12. S. J. Chung, S. M. Armasu, J. M. Biernacka, K. J. Anderson, T. G. Lesnick, D. N. Rider, J. M. Cunningham, J. Eric Ahlskog, R. Frigerio, D. M. Maraganore, Genomic determinants of motor and cognitive outcomes in Parkinson’s disease. Parkinsonism Relat Disord 18, 881–886 (2012).

13. M. A. Nalls, C. Blauwendraat, C. L. Vallerga, K. Heilbron, S. Bandres-Ciga, D. Chang, M. Tan, D. A. Kia, A. J. Noyce, A. Xue, J. Bras, E. Young, R. von Coelln, J. Simon-Sanchez, C. Schulte, M. Sharma, L. Krohn, L. Pihlstrom, A. Siitonen, H. Iwaki, H. Leonard, F. Faghri, J. R. Gibbs, D. G. Hernandez, S. W. Scholz, J. A. Botia, M. Martinez, J. C. Corvol, S. Lesage, J. Jankovic, L. M. Shulman, M. Sutherland, P. Tienari, K. Majamaa, M. Toft, O. A. Andreassen, T. Bangale, A. Brice, J. Yang, Z. Gan-Or, T. Gasser, P. Heutink, J. M. Shulman, N. W. Wood, D. A. Hinds, J. A. Hardy, H. R. Morris, J. Gratten, P. M. Visscher, R. R. Graham, A. B. Singleton, T. andMe Research, C. System Genomics of Parkinson’s Disease, C. International Parkinson’s Disease Genomics, Identification of novel risk loci, causal insights, and heritable risk for Parkinson’s disease: a meta-analysis of genome-wide association studies. Lancet Neurol 18, 1091–1102 (2019).

14. C. Blauwendraat, M. A. Nalls, A. B. Singleton, The genetic architecture of Parkinson’s disease. Lancet Neurol 19, 170–178 (2020).

15. D. G. Hernandez, X. Reed, A. B. Singleton, Genetics in Parkinson disease: Mendelian versus non-Mendelian inheritance. J Neurochem 139 Suppl 1, 59–74 (2016).

16. K. R. Kumar, A. Djarmati-Westenberger, A. Grunewald, Genetics of Parkinson’s disease. Semin Neurol 31, 433–440 (2011).

17. N. V. N. Carullo, J. J. Day, Genomic Enhancers in Brain Health and Disease. Genes (Basel) 10, (2019).

18. J. M. Karnuta, P. C. Scacheri, Enhancers: bridging the gap between gene control and human disease. Hum Mol Genet 27, R219–R227 (2018).

19. A. R. Niederriter, A. Varshney, S. C. Parker, D. M. Martin, Super Enhancers in Cancers, Complex Disease, and Developmental Disorders. Genes (Basel) 6, 1183–1200 (2015).

20. I. Jung, A. Schmitt, Y. Diao, A. J. Lee, T. Liu, D. Yang, C. Tan, J. Eom, M. Chan, S. Chee, Z. Chiang, C. Kim, E. Masliah, C. L. Barr, B. Li, S. Kuan, D. Kim, B. Ren, A compendium of promoter-centered long-range chromatin interactions in the human genome. Nat Genet 51, 1442–1449 (2019).

21. M. T. Maurano, R. Humbert, E. Rynes, R. E. Thurman, E. Haugen, H. Wang, A. P. Reynolds, R. Sandstrom, H. Qu, J. Brody, A. Shafer, F. Neri, K. Lee, T. Kutyavin, S. Stehling-Sun, A. K. Johnson, T. K. Canfield, E. Giste, M. Diegel, D. Bates, R. S. Hansen, S. Neph, P. J. Sabo, S. Heimfeld, A. Raubitschek, S. Ziegler, C. Cotsapas, N. Sotoodehnia, I. Glass, S. R. Sunyaev, R. Kaul, J. A. Stamatoyannopoulos, Systematic localization of common disease-associated variation in regulatory DNA. Science 337, 1190–1195 (2012).

22. L. Toker, G. T. Tran, J. Sundaresan, O. B. Tysnes, G. Alves, K. Haugarvoll, G. S. Nido, C. Dolle, C. Tzoulis, Genome-wide histone acetylation analysis reveals altered transcriptional regulation in the Parkinson’s disease brain. Mol Neurodegener 16, 31 (2021).

23. M. R. Corces, A. Shcherbina, S. Kundu, M. J. Gloudemans, L. Fresard, J. M. Granja, B. H. Louie, T. Eulalio, S. Shams, S. T. Bagdatli, M. R. Mumbach, B. Liu, K. S. Montine, W. J. Greenleaf, A. Kundaje, S. B. Montgomery, H. Y. Chang, T. J. Montine, Single-cell epigenomic analyses implicate candidate causal variants at inherited risk loci for Alzheimer’s and Parkinson’s diseases. Nat Genet 52, 1158–1168 (2020).

24. S. Morabito, E. Miyoshi, N. Michael, S. Shahin, A. C. Martini, E. Head, J. Silva, K. Leavy, M. Perez-Rosendahl, V. Swarup, Single-nucleus chromatin accessibility and transcriptomic characterization of Alzheimer’s disease. Nat Genet 53, 1143–1155 (2021).

25. M. J. Fullwood, M. H. Liu, Y. F. Pan, J. Liu, H. Xu, Y. B. Mohamed, Y. L. Orlov, S. Velkov, A. Ho, P. H. Mei, E. G. Chew, P. Y. Huang, W. J. Welboren, Y. Han, H. S. Ooi, P. N. Ariyaratne, V. B. Vega, Y. Luo, P. Y. Tan, P. Y. Choy, K. D. Wansa, B. Zhao, K. S. Lim, S. C. Leow, J. S. Yow, R. Joseph, H. Li, K. V. Desai, J. S. Thomsen, Y. K. Lee, R. K. Karuturi, T. Herve, G. Bourque, H. G. Stunnenberg, X. Ruan, V. Cacheux-Rataboul, W. K. Sung, E. T. Liu, C. L. Wei, E. Cheung, Y. Ruan, An oestrogen-receptor-alpha-bound human chromatin interactome. Nature 462, 58–64 (2009).

26. E. Lieberman-Aiden, N. L. van Berkum, L. Williams, M. Imakaev, T. Ragoczy, A. Telling, I. Amit, B. R. Lajoie, P. J. Sabo, M. O. Dorschner, R. Sandstrom, B. Bernstein, M. A. Bender, M. Groudine, A. Gnirke, J. Stamatoyannopoulos, L. A. Mirny, E. S. Lander, J. Dekker, Comprehensive mapping of long-range interactions reveals folding principles of the human genome. Science 326, 289–293 (2009).

27. M. R. Mumbach, A. J. Rubin, R. A. Flynn, C. Dai, P. A. Khavari, W. J. Greenleaf, H. Y. Chang, HiChIP: efficient and sensitive analysis of protein-directed genome architecture. Nat Methods 13, 919–922 (2016).

28. R. Fang, M. Yu, G. Li, S. Chee, T. Liu, A. D. Schmitt, B. Ren, Mapping of long-range chromatin interactions by proximity ligation-assisted ChIP-seq. Cell Res 26, 1345–1348 (2016).

29. B. Mifsud, F. Tavares-Cadete, A. N. Young, R. Sugar, S. Schoenfelder, L. Ferreira, S. W. Wingett, S. Andrews, W. Grey, P. A. Ewels, B. Herman, S. Happe, A. Higgs, E. LeProust, G. A. Follows, P. Fraser, N. M. Luscombe, C. S. Osborne, Mapping long-range promoter contacts in human cells with high-resolution capture Hi-C. Nat Genet 47, 598–606 (2015).

30. A. Sanyal, B. R. Lajoie, G. Jain, J. Dekker, The long-range interaction landscape of gene promoters. Nature 489, 109–113 (2012).

31. J. R. Dixon, I. Jung, S. Selvaraj, Y. Shen, J. E. Antosiewicz-Bourget, A. Y. Lee, Z. Ye, A. Kim, N. Rajagopal, W. Xie, Y. Diao, J. Liang, H. Zhao, V. V. Lobanenkov, J. R. Ecker, J. A. Thomson, B. Ren, Chromatin architecture reorganization during stem cell differentiation. Nature 518, 331–336 (2015).

32. F. Jin, Y. Li, J. R. Dixon, S. Selvaraj, Z. Ye, A. Y. Lee, C. A. Yen, A. D. Schmitt, C. A. Espinoza, B. Ren, A high-resolution map of the three-dimensional chromatin interactome in human cells. Nature 503, 290–294 (2013).

33. S. S. Rao, M. H. Huntley, N. C. Durand, E. K. Stamenova, I. D. Bochkov, J. T. Robinson, A. L. Sanborn, I. Machol, A. D. Omer, E. S. Lander, E. L. Aiden, A 3D map of the human genome at kilobase resolution reveals principles of chromatin looping. Cell 159, 1665–1680 (2014).

34. A. Nott, I. R. Holtman, N. G. Coufal, J. C. M. Schlachetzki, M. Yu, R. Hu, C. Z. Han, M. Pena, J. Xiao, Y. Wu, Z. Keulen, M. P. Pasillas, C. O’Connor, C. K. Nickl, S. T. Schafer, Z. Shen, R. A. Rissman, J. B. Brewer, D. Gosselin, D. D. Gonda, M. L. Levy, M. G. Rosenfeld, G. McVicker, F. H. Gage, B. Ren, C. K. Glass, Brain cell type-specific enhancer-promoter interactome maps and disease-risk association. Science 366, 1134–1139 (2019).

35. D. Agarwal, C. Sandor, V. Volpato, T. M. Caffrey, J. Monzon-Sandoval, R. Bowden, J. Alegre-Abarrategui, R. Wade-Martins, C. Webber, A single-cell atlas of the human substantia nigra reveals cell-specific pathways associated with neurological disorders. Nat Commun 11, 4183 (2020).

36. X. L. Gu, C. X. Long, L. Sun, C. Xie, X. Lin, H. Cai, Astrocytic expression of Parkinson’s disease-related A53T alpha-synuclein causes neurodegeneration in mice. Mol Brain 3, 12 (2010).

37. M. E. Hamby, M. V. Sofroniew, Reactive astrocytes as therapeutic targets for CNS disorders. Neurotherapeutics 7, 494–506 (2010).

38. K. J. Billingsley, S. Bandres-Ciga, S. Saez-Atienzar, A. B. Singleton, Genetic risk factors in Parkinson’s disease. Cell Tissue Res 373, 9–20 (2018).

39. M. D. Robinson, D. J. McCarthy, G. K. Smyth, edgeR: a Bioconductor package for differential expression analysis of digital gene expression data. Bioinformatics 26, 139–140 (2010).

40. C. P. Fulco, J. Nasser, T. R. Jones, G. Munson, D. T. Bergman, V. Subramanian, S. R. Grossman, R. Anyoha, B. R. Doughty, T. A. Patwardhan, T. H. Nguyen, M. Kane, E. M. Perez, N. C. Durand, C. A. Lareau, E. K. Stamenova, E. L. Aiden, E. S. Lander, J. M. Engreitz, Activity-by-contact model of enhancer-promoter regulation from thousands of CRISPR perturbations. Nat Genet 51, 1664–1669 (2019).

41. F. Ay, T. L. Bailey, W. S. Noble, Statistical confidence estimation for Hi-C data reveals regulatory chromatin contacts. Genome Res 24, 999–1011 (2014).

42. M. Claussnitzer, S. N. Dankel, K. H. Kim, G. Quon, W. Meuleman, C. Haugen, V. Glunk, I. S. Sousa, J. L. Beaudry, V. Puviindran, N. A. Abdennur, J. Liu, P. A. Svensson, Y. H. Hsu, D. J. Drucker, G. Mellgren, C. C. Hui, H. Hauner, M. Kellis, FTO Obesity Variant Circuitry and Adipocyte Browning in Humans. N Engl J Med 373, 895–907 (2015).

43. B. K. Bulik-Sullivan, P. R. Loh, H. K. Finucane, S. Ripke, J. Yang, C. Schizophrenia Working Group of the Psychiatric Genomics, N. Patterson, M. J. Daly, A. L. Price, B. M. Neale, LD Score regression distinguishes confounding from polygenicity in genome-wide association studies. Nat Genet 47, 291–295 (2015).

44. H. K. Finucane, Y. A. Reshef, V. Anttila, K. Slowikowski, A. Gusev, A. Byrnes, S. Gazal, P. R. Loh, C. Lareau, N. Shoresh, G. Genovese, A. Saunders, E. Macosko, S. Pollack, C. Brainstorm, J. R. B. Perry, J. D. Buenrostro, B. E. Bernstein, S. Raychaudhuri, S. McCarroll, B. M. Neale, A. L. Price, Heritability enrichment of specifically expressed genes identifies disease-relevant tissues and cell types. Nat Genet 50, 621–629 (2018).

45. I. E. Jansen, J. E. Savage, K. Watanabe, J. Bryois, D. M. Williams, S. Steinberg, J. Sealock, I. K. Karlsson, S. Hagg, L. Athanasiu, N. Voyle, P. Proitsi, A. Witoelar, S. Stringer, D. Aarsland, I. S. Almdahl, F. Andersen, S. Bergh, F. Bettella, S. Bjornsson, A. Braekhus, G. Brathen, C. de Leeuw, R. S. Desikan, S. Djurovic, L. Dumitrescu, T. Fladby, T. J. Hohman, P. V. Jonsson, S. J. Kiddle, A. Rongve, I. Saltvedt, S. B. Sando, G. Selbaek, M. Shoai, N. G. Skene, J. Snaedal, E. Stordal, I. D. Ulstein, Y. Wang, L. R. White, J. Hardy, J. Hjerling-Leffler, P. F. Sullivan, W. M. van der Flier, R. Dobson, L. K. Davis, H. Stefansson, K. Stefansson, N. L. Pedersen, S. Ripke, O. A. Andreassen, D. Posthuma, Genome-wide meta-analysis identifies new loci and functional pathways influencing Alzheimer’s disease risk. Nat Genet 51, 404–413 (2019).

46. B. W. Kunkle, B. Grenier-Boley, R. Sims, J. C. Bis, V. Damotte, A. C. Naj, A. Boland, M. Vronskaya, S. J. van der Lee, A. Amlie-Wolf, C. Bellenguez, A. Frizatti, V. Chouraki, E. R. Martin, K. Sleegers, N. Badarinarayan, J. Jakobsdottir, K. L. Hamilton-Nelson, S. Moreno-Grau, R. Olaso, R. Raybould, Y. Chen, A. B. Kuzma, M. Hiltunen, T. Morgan, S. Ahmad, B. N. Vardarajan, J. Epelbaum, P. Hoffmann, M. Boada, G. W. Beecham, J. G. Garnier, D. Harold, A. L. Fitzpatrick, O. Valladares, M. L. Moutet, A. Gerrish, A. V. Smith, L. Qu, D. Bacq, N. Denning, X. Jian, Y. Zhao, M. Del Zompo, N. C. Fox, S. H. Choi, I. Mateo, J. T. Hughes, H. H. Adams, J. Malamon, F. Sanchez-Garcia, Y. Patel, J. A. Brody, B. A. Dombroski, M. C. D. Naranjo, M. Daniilidou, G. Eiriksdottir, S. Mukherjee, D. Wallon, J. Uphill, T. Aspelund, L. B. Cantwell, F. Garzia, D. Galimberti, E. Hofer, M. Butkiewicz, B. Fin, E. Scarpini, C. Sarnowski, W. S. Bush, S. Meslage, J. Kornhuber, C. C. White, Y. Song, R. C. Barber, S. Engelborghs, S. Sordon, D. Voijnovic, P. M. Adams, R. Vandenberghe, M. Mayhaus, L. A. Cupples, M. S. Albert, P. P. De Deyn, W. Gu, J. J. Himali, D. Beekly, A. Squassina, A. M. Hartmann, A. Orellana, D. Blacker, E. Rodriguez-Rodriguez, S. Lovestone, M. E. Garcia, R. S. Doody, C. Munoz-Fernadez, R. Sussams, H. Lin, T. J. Fairchild, Y. A. Benito, C. Holmes, H. Karamujic-Comic, M. P. Frosch, H. Thonberg, W. Maier, G. Roshchupkin, B. Ghetti, V. Giedraitis, A. Kawalia, S. Li, R. M. Huebinger, L. Kilander, S. Moebus, I. Hernandez, M. I. Kamboh, R. Brundin, J. Turton, Q. Yang, M. J. Katz, L. Concari, J. Lord, A. S. Beiser, C. D. Keene, S. Helisalmi, I. Kloszewska, W. A. Kukull, A. M. Koivisto, A. Lynch, L. Tarraga, E. B. Larson, A. Haapasalo, B. Lawlor, T. H. Mosley, R. B. Lipton, V. Solfrizzi, M. Gill, W. T. Longstreth, Jr., T. J. Montine, V. Frisardi, M. Diez-Fairen, F. Rivadeneira, R. C. Petersen, V. Deramecourt, I. Alvarez, F. Salani, A. Ciaramella, E. Boerwinkle, E. M. Reiman, N. Fievet, J. I. Rotter, J. S. Reisch, O. Hanon, C. Cupidi, A. G. Andre Uitterlinden, D. R. Royall, C. Dufouil, R. G. Maletta, I. de Rojas, M. Sano, A. Brice, R. Cecchetti, P. S. George-Hyslop, K. Ritchie, M. Tsolaki, D. W. Tsuang, B. Dubois, D. Craig, C. K. Wu, H. Soininen, D. Avramidou, R. L. Albin, L. Fratiglioni, A. Germanou, L. G. Apostolova, L. Keller, M. Koutroumani, S. E. Arnold, F. Panza, O. Gkatzima, S. Asthana, D. Hannequin, P. Whitehead, C. S. Atwood, P. Caffarra, H. Hampel, I. Quintela, A. Carracedo, L. Lannfelt, D. C. Rubinsztein, L. L. Barnes, F. Pasquier, L. Frolich, S. Barral, B. McGuinness, T. G. Beach, J. A. Johnston, J. T. Becker, P. Passmore, E. H. Bigio, J. M. Schott, T. D. Bird, J. D. Warren, B. F. Boeve, M. K. Lupton, J. D. Bowen, P. Proitsi, A. Boxer, J. F. Powell, J. R. Burke, J. S. K. Kauwe, J. M. Burns, M. Mancuso, J. D. Buxbaum, U. Bonuccelli, N. J. Cairns, A. McQuillin, C. Cao, G. Livingston, C. S. Carlson, N. J. Bass, C. M. Carlsson, J. Hardy, R. M. Carney, J. Bras, M. M. Carrasquillo, R. Guerreiro, M. Allen, H. C. Chui, E. Fisher, C. Masullo, E. A. Crocco, C. DeCarli, G. Bisceglio, M. Dick, L. Ma, R. Duara, N. R. Graff-Radford, D. A. Evans, A. Hodges, K. M. Faber, M. Scherer, K. B. Fallon, M. Riemenschneider, D. W. Fardo, R. Heun, M. R. Farlow, H. Kolsch, S. Ferris, M. Leber, T. M. Foroud, I. Heuser, D. R. Galasko, I. Giegling, M. Gearing, M. Hull, D. H. Geschwind, J. R. Gilbert, J. Morris, R. C. Green, K. Mayo, J. H. Growdon, T. Feulner, R. L. Hamilton, L. E. Harrell, D. Drichel, L. S. Honig, T. D. Cushion, M. J. Huentelman, P. Hollingworth, C. M. Hulette, B. T. Hyman, R. Marshall, G. P. Jarvik, A. Meggy, E. Abner, G. E. Menzies, L. W. Jin, G. Leonenko, L. M. Real, G. R. Jun, C. T. Baldwin, D. Grozeva, A. Karydas, G. Russo, J. A. Kaye, R. Kim, F. Jessen, N. W. Kowall, B. Vellas, J. H. Kramer, E. Vardy, F. M. LaFerla, K. H. Jockel, J. J. Lah, M. Dichgans, J. B. Leverenz, D. Mann, A. I. Levey, S. Pickering-Brown, A. P. Lieberman, N. Klopp, K. L. Lunetta, H. E. Wichmann, C. G. Lyketsos, K. Morgan, D. C. Marson, K. Brown, F. Martiniuk, C. Medway, D. C. Mash, M. M. Nothen, E. Masliah, N. M. Hooper, W. C. McCormick, A. Daniele, S. M. McCurry, A. Bayer, A. N. McDavid, J. Gallacher, A. C. McKee, H. van den Bussche, M. Mesulam, C. Brayne, B. L. Miller, S. Riedel-Heller, C. A. Miller, J. W. Miller, A. Al-Chalabi, J. C. Morris, C. E. Shaw, A. J. Myers, J. Wiltfang, S. O’Bryant, J. M. Olichney, V. Alvarez, J. E. Parisi, A. B. Singleton, H. L. Paulson, J. Collinge, W. R. Perry, S. Mead, E. Peskind, D. H. Cribbs, M. Rossor, A. Pierce, N. S. Ryan, W. W. Poon, B. Nacmias, H. Potter, S. Sorbi, J. F. Quinn, E. Sacchinelli, A. Raj, G. Spalletta, M. Raskind, C. Caltagirone, P. Bossu, M. D. Orfei, B. Reisberg, R. Clarke, C. Reitz, A. D. Smith, J. M. Ringman, D. Warden, E. D. Roberson, G. Wilcock, E. Rogaeva, A. C. Bruni, H. J. Rosen, M. Gallo, R. N. Rosenberg, Y. Ben-Shlomo, M. A. Sager, P. Mecocci, A. J. Saykin, P. Pastor, M. L. Cuccaro, J. M. Vance, J. A. Schneider, L. S. Schneider, S. Slifer, W. W. Seeley, A. G. Smith, J. A. Sonnen, S. Spina, R. A. Stern, R. H. Swerdlow, M. Tang, R. E. Tanzi, J. Q. Trojanowski, J. C. Troncoso, V. M. Van Deerlin, L. J. Van Eldik, H. V. Vinters, J. P. Vonsattel, S. Weintraub, K. A. Welsh-Bohmer, K. C. Wilhelmsen, J. Williamson, T. S. Wingo, R. L. Woltjer, C. B. Wright, C. E. Yu, L. Yu, Y. Saba, A. Pilotto, M. J. Bullido, O. Peters, P. K. Crane, D. Bennett, P. Bosco, E. Coto, V. Boccardi, P. L. De Jager, A. Lleo, N. Warner, O. L. Lopez, M. Ingelsson, P. Deloukas, C. Cruchaga, C. Graff, R. Gwilliam, M. Fornage, A. M. Goate, P. Sanchez-Juan, P. G. Kehoe, N. Amin, N. Ertekin-Taner, C. Berr, S. Debette, S. Love, L. J. Launer, S. G. Younkin, J. F. Dartigues, C. Corcoran, M. A. Ikram, D. W. Dickson, G. Nicolas, D. Campion, J. Tschanz, H. Schmidt, H. Hakonarson, J. Clarimon, R. Munger, R. Schmidt, L. A. Farrer, C. Van Broeckhoven, C. O. D. M, A. L. DeStefano, L. Jones, J. L. Haines, J. F. Deleuze, M. J. Owen, V. Gudnason, R. Mayeux, V. Escott-Price, B. M. Psaty, A. Ramirez, L. S. Wang, A. Ruiz, C. M. van Duijn, P. A. Holmans, S. Seshadri, J. Williams, P. Amouyel, G. D. Schellenberg, J. C. Lambert, M. A. Pericak-Vance, C. Alzheimer Disease Genetics, I. European Alzheimer’s Disease, H. Cohorts for, C. Aging Research in Genomic Epidemiology, Genetic, P. Environmental Risk in Ad/Defining Genetic, C. Environmental Risk for Alzheimer’s Disease, Genetic meta-analysis of diagnosed Alzheimer’s disease identifies new risk loci and implicates Abeta, tau, immunity and lipid processing. Nat Genet 51, 414–430 (2019).

47. W. van Rheenen, A. Shatunov, A. M. Dekker, R. L. McLaughlin, F. P. Diekstra, S. L. Pulit, R. A. van der Spek, U. Vosa, S. de Jong, M. R. Robinson, J. Yang, I. Fogh, P. T. van Doormaal, G. H. Tazelaar, M. Koppers, A. M. Blokhuis, W. Sproviero, A. R. Jones, K. P. Kenna, K. R. van Eijk, O. Harschnitz, R. D. Schellevis, W. J. Brands, J. Medic, A. Menelaou, A. Vajda, N. Ticozzi, K. Lin, B. Rogelj, K. Vrabec, M. Ravnik-Glavac, B. Koritnik, J. Zidar, L. Leonardis, L. D. Groselj, S. Millecamps, F. Salachas, V. Meininger, M. de Carvalho, S. Pinto, J. S. Mora, R. Rojas-Garcia, M. Polak, S. Chandran, S. Colville, R. Swingler, K. E. Morrison, P. J. Shaw, J. Hardy, R. W. Orrell, A. Pittman, K. Sidle, P. Fratta, A. Malaspina, S. Topp, S. Petri, S. Abdulla, C. Drepper, M. Sendtner, T. Meyer, R. A. Ophoff, K. A. Staats, M. Wiedau-Pazos, C. Lomen-Hoerth, V. M. Van Deerlin, J. Q. Trojanowski, L. Elman, L. McCluskey, A. N. Basak, C. Tunca, H. Hamzeiy, Y. Parman, T. Meitinger, P. Lichtner, M. Radivojkov-Blagojevic, C. R. Andres, C. Maurel, G. Bensimon, B. Landwehrmeyer, A. Brice, C. A. Payan, S. Saker-Delye, A. Durr, N. W. Wood, L. Tittmann, W. Lieb, A. Franke, M. Rietschel, S. Cichon, M. M. Nothen, P. Amouyel, C. Tzourio, J. F. Dartigues, A. G. Uitterlinden, F. Rivadeneira, K. Estrada, A. Hofman, C. Curtis, H. M. Blauw, A. J. van der Kooi, M. de Visser, A. Goris, M. Weber, C. E. Shaw, B. N. Smith, O. Pansarasa, C. Cereda, R. Del Bo, G. P. Comi, S. D’Alfonso, C. Bertolin, G. Soraru, L. Mazzini, V. Pensato, C. Gellera, C. Tiloca, A. Ratti, A. Calvo, C. Moglia, M. Brunetti, S. Arcuti, R. Capozzo, C. Zecca, C. Lunetta, S. Penco, N. Riva, A. Padovani, M. Filosto, B. Muller, R. J. Stuit, P. Registry, S. Group, S. Registry, F. S. Consortium, S. Consortium, N. S. Group, I. Blair, K. Zhang, E. P. McCann, J. A. Fifita, G. A. Nicholson, D. B. Rowe, R. Pamphlett, M. C. Kiernan, J. Grosskreutz, O. W. Witte, T. Ringer, T. Prell, B. Stubendorff, I. Kurth, C. A. Hubner, P. N. Leigh, F. Casale, A. Chio, E. Beghi, E. Pupillo, R. Tortelli, G. Logroscino, J. Powell, A. C. Ludolph, J. H. Weishaupt, W. Robberecht, P. Van Damme, L. Franke, T. H. Pers, R. H. Brown, J. D. Glass, J. E. Landers, O. Hardiman, P. M. Andersen, P. Corcia, P. Vourc’h, V. Silani, N. R. Wray, P. M. Visscher, P. I. de Bakker, M. A. van Es, R. J. Pasterkamp, C. M. Lewis, G. Breen, A. Al-Chalabi, L. H. van den Berg, J. H. Veldink, Genome-wide association analyses identify new risk variants and the genetic architecture of amyotrophic lateral sclerosis. Nat Genet 48, 1043–1048 (2016).

48. J. Grove, S. Ripke, T. D. Als, M. Mattheisen, R. K. Walters, H. Won, J. Pallesen, E. Agerbo, O. A. Andreassen, R. Anney, S. Awashti, R. Belliveau, F. Bettella, J. D. Buxbaum, J. Bybjerg-Grauholm, M. Baekvad-Hansen, F. Cerrato, K. Chambert, J. H. Christensen, C. Churchhouse, K. Dellenvall, D. Demontis, S. De Rubeis, B. Devlin, S. Djurovic, A. L. Dumont, J. I. Goldstein, C. S. Hansen, M. E. Hauberg, M. V. Hollegaard, S. Hope, D. P. Howrigan, H. Huang, C. M. Hultman, L. Klei, J. Maller, J. Martin, A. R. Martin, J. L. Moran, M. Nyegaard, T. Naerland, D. S. Palmer, A. Palotie, C. B. Pedersen, M. G. Pedersen, T. dPoterba, J. B. Poulsen, B. S. Pourcain, P. Qvist, K. Rehnstrom, A. Reichenberg, J. Reichert, E. B. Robinson, K. Roeder, P. Roussos, E. Saemundsen, S. Sandin, F. K. Satterstrom, G. Davey Smith, H. Stefansson, S. Steinberg, C. R. Stevens, P. F. Sullivan, P. Turley, G. B. Walters, X. Xu, C. Autism Spectrum Disorder Working Group of the Psychiatric Genomics, Bupgen, C. Major Depressive Disorder Working Group of the Psychiatric Genomics, andMe Research, K. Stefansson, D. H. Geschwind, M. Nordentoft, D. M. Hougaard, T. Werge, O. Mors, P. B. Mortensen, B. M. Neale, M. J. Daly, A. D. Borglum, Identification of common genetic risk variants for autism spectrum disorder. Nat Genet 51, 431–444 (2019).

49. A. F. Pardinas, P. Holmans, A. J. Pocklington, V. Escott-Price, S. Ripke, N. Carrera, S. E. Legge, S. Bishop, D. Cameron, M. L. Hamshere, J. Han, L. Hubbard, A. Lynham, K. Mantripragada, E. Rees, J. H. MacCabe, S. A. McCarroll, B. T. Baune, G. Breen, E. M. Byrne, U. Dannlowski, T. C. Eley, C. Hayward, N. G. Martin, A. M. McIntosh, R. Plomin, D. J. Porteous, N. R. Wray, A. Caballero, D. H. Geschwind, L. M. Huckins, D. M. Ruderfer, E. Santiago, P. Sklar, E. A. Stahl, H. Won, E. Agerbo, T. D. Als, O. A. Andreassen, M. Baekvad-Hansen, P. B. Mortensen, C. B. Pedersen, A. D. Borglum, J. Bybjerg-Grauholm, S. Djurovic, N. Durmishi, M. G. Pedersen, V. Golimbet, J. Grove, D. M. Hougaard, M. Mattheisen, E. Molden, O. Mors, M. Nordentoft, M. Pejovic-Milovancevic, E. Sigurdsson, T. Silagadze, C. S. Hansen, K. Stefansson, H. Stefansson, S. Steinberg, S. Tosato, T. Werge, G. Consortium, C. Consortium, D. A. Collier, D. Rujescu, G. Kirov, M. J. Owen, M. C. O’Donovan, J. T. R. Walters, Common schizophrenia alleles are enriched in mutation-intolerant genes and in regions under strong background selection. Nat Genet 50, 381–389 (2018).

50. L. Gan, M. R. Cookson, L. Petrucelli, A. R. La Spada, Converging pathways in neurodegeneration, from genetics to mechanisms. Nat Neurosci 21, 1300–1309 (2018).

51. A. Elkouzi, V. Vedam-Mai, R. S. Eisinger, M. S. Okun, Emerging therapies in Parkinson disease - repurposed drugs and new approaches. Nat Rev Neurol 15, 204–223 (2019).

52. I. Alecu, S. A. L. Bennett, Dysregulated Lipid Metabolism and Its Role in alpha-Synucleinopathy in Parkinson’s Disease. Front Neurosci 13, 328 (2019).

53. E. C. Hirsch, S. Hunot, Neuroinflammation in Parkinson’s disease: a target for neuroprotection? Lancet Neurol 8, 382–397 (2009).

54. F. Hopfner, E. C. Schulte, B. Mollenhauer, B. Bereznai, F. Knauf, P. Lichtner, A. Zimprich, D. Haubenberger, W. Pirker, T. Brucke, A. Peters, C. Gieger, G. Kuhlenbaumer, C. Trenkwalder, J. Winkelmann, The role of SCARB2 as susceptibility factor in Parkinson’s disease. Mov Disord 28, 538–540 (2013).

55. H. Michelakakis, G. Xiromerisiou, E. Dardiotis, M. Bozi, D. Vassilatis, P. M. Kountra, G. Patramani, M. Moraitou, D. Papadimitriou, E. Stamboulis, L. Stefanis, E. Zintzaras, G. M. Hadjigeorgiou, Evidence of an association between the scavenger receptor class B member 2 gene and Parkinson’s disease. Mov Disord 27, 400–405 (2012).

56. C. Blauwendraat, K. Heilbron, C. L. Vallerga, S. Bandres-Ciga, R. von Coelln, L. Pihlstrom, J. Simon-Sanchez, C. Schulte, M. Sharma, L. Krohn, A. Siitonen, H. Iwaki, H. Leonard, A. J. Noyce, M. Tan, J. R. Gibbs, D. G. Hernandez, S. W. Scholz, J. Jankovic, L. M. Shulman, S. Lesage, J. C. Corvol, A. Brice, J. J. van Hilten, J. Marinus, T. andMe Research, J. Eerola-Rautio, P. Tienari, K. Majamaa, M. Toft, D. G. Grosset, T. Gasser, P. Heutink, J. M. Shulman, N. Wood, J. Hardy, H. R. Morris, D. A. Hinds, J. Gratten, P. M. Visscher, Z. Gan-Or, M. A. Nalls, A. B. Singleton, C. International Parkinson’s Disease Genomics, Parkinson’s disease age at onset genome-wide association study: Defining heritability, genetic loci, and alpha-synuclein mechanisms. Mov Disord 34, 866–875 (2019).

57. J. Bove, M. Martinez-Vicente, B. Dehay, C. Perier, A. Recasens, A. Bombrun, B. Antonsson, M. Vila, BAX channel activity mediates lysosomal disruption linked to Parkinson disease. Autophagy 10, 889–900 (2014).

58. D. M. Arduino, A. R. Esteves, S. M. Cardoso, Mitochondria drive autophagy pathology via microtubule disassembly: a new hypothesis for Parkinson disease. Autophagy 9, 112–114 (2013).

59. J. Obergasteiger, G. Frapporti, P. P. Pramstaller, A. A. Hicks, M. Volta, A new hypothesis for Parkinson’s disease pathogenesis: GTPase-p38 MAPK signaling and autophagy as convergence points of etiology and genomics. Mol Neurodegener 13, 40 (2018).

60. T. Nagano, Y. Lubling, C. Varnai, C. Dudley, W. Leung, Y. Baran, N. Mendelson Cohen, S. Wingett, P. Fraser, A. Tanay, Cell-cycle dynamics of chromosomal organization at single-cell resolution. Nature 547, 61–67 (2017).

61. G. Li, Y. Liu, Y. Zhang, N. Kubo, M. Yu, R. Fang, M. Kellis, B. Ren, Joint profiling of DNA methylation and chromatin architecture in single cells. Nat Methods 16, 991–993 (2019).

62. D. S. Lee, C. Luo, J. Zhou, S. Chandran, A. Rivkin, A. Bartlett, J. R. Nery, C. Fitzpatrick, C. O’Connor, J. R. Dixon, J. R. Ecker, Simultaneous profiling of 3D genome structure and DNA methylation in single human cells. Nat Methods 16, 999–1006 (2019).

63. J. S. Schweitzer, B. Song, T. M. Herrington, T. Y. Park, N. Lee, S. Ko, J. Jeon, Y. Cha, K. Kim, Q. Li, C. Henchcliffe, M. Kaplitt, C. Neff, O. Rapalino, H. Seo, I. H. Lee, J. Kim, T. Kim, G. A. Petsko, J. Ritz, B. M. Cohen, S. W. Kong, P. Leblanc, B. S. Carter, K. S. Kim, Personalized iPSC-Derived Dopamine Progenitor Cells for Parkinson’s Disease. N Engl J Med 382, 1926–1932 (2020).

64. H. Braak, E. Braak, Neuropathological stageing of Alzheimer-related changes. Acta Neuropathol 82, 239–259 (1991).

65. H. Heaton, A. M. Talman, A. Knights, M. Imaz, D. J. Gaffney, R. Durbin, M. Hemberg, M. K. N. Lawniczak, Souporcell: robust clustering of single-cell RNA-seq data by genotype without reference genotypes. Nat Methods 17, 615–620 (2020).

66. G. X. Zheng, J. M. Terry, P. Belgrader, P. Ryvkin, Z. W. Bent, R. Wilson, S. B. Ziraldo, T. D. Wheeler, G. P. McDermott, J. Zhu, M. T. Gregory, J. Shuga, L. Montesclaros, J. G. Underwood, D. A. Masquelier, S. Y. Nishimura, M. Schnall-Levin, P. W. Wyatt, C. M. Hindson, R. Bharadwaj, A. Wong, K. D. Ness, L. W. Beppu, H. J. Deeg, C. McFarland, K. R. Loeb, W. J. Valente, N. G. Ericson, E. A. Stevens, J. P. Radich, T. S. Mikkelsen, B. J. Hindson, J. H. Bielas, Massively parallel digital transcriptional profiling of single cells. Nat Commun 8, 14049 (2017).

67. Y. Hao, S. Hao, E. Andersen-Nissen, W. M. Mauck, 3rd, S. Zheng, A. Butler, M. J. Lee, A. J. Wilk, C. Darby, M. Zager, P. Hoffman, M. Stoeckius, E. Papalexi, E. P. Mimitou, J. Jain, A. Srivastava, T. Stuart, L. M. Fleming, B. Yeung, A. J. Rogers, J. M. McElrath, C. A. Blish, R. Gottardo, P. Smibert, R. Satija, Integrated analysis of multimodal single-cell data. Cell 184, 3573–3587 e3529 (2021).

68. S. L. Wolock, R. Lopez, A. M. Klein, Scrublet: Computational Identification of Cell Doublets in Single-Cell Transcriptomic Data. Cell Syst 8, 281–291 e289 (2019).

69. I. Korsunsky, N. Millard, J. Fan, K. Slowikowski, F. Zhang, K. Wei, Y. Baglaenko, M. Brenner, P. R. Loh, S. Raychaudhuri, Fast, sensitive and accurate integration of single-cell data with Harmony. Nat Methods 16, 1289–1296 (2019).

70. D. van Dijk, R. Sharma, J. Nainys, K. Yim, P. Kathail, A. J. Carr, C. Burdziak, K. R. Moon, C. L. Chaffer, D. Pattabiraman, B. Bierie, L. Mazutis, G. Wolf, S. Krishnaswamy, D. Pe’er, Recovering Gene Interactions from Single-Cell Data Using Data Diffusion. Cell 174, 716–729 e727 (2018).

71. A. T. Satpathy, J. M. Granja, K. E. Yost, Y. Qi, F. Meschi, G. P. McDermott, B. N. Olsen, M. R. Mumbach, S. E. Pierce, M. R. Corces, P. Shah, J. C. Bell, D. Jhutty, C. M. Nemec, J. Wang, L. Wang, Y. Yin, P. G. Giresi, A. L. S. Chang, G. X. Y. Zheng, W. J. Greenleaf, H. Y. Chang, Massively parallel single-cell chromatin landscapes of human immune cell development and intratumoral T cell exhaustion. Nat Biotechnol 37, 925–936 (2019).

72. T. Stuart, A. Srivastava, S. Madad, C. A. Lareau, R. Satija, Single-cell chromatin state analysis with Signac. Nat Methods 18, 1333–1341 (2021).

73. Y. Zhang, T. Liu, C. A. Meyer, J. Eeckhoute, D. S. Johnson, B. E. Bernstein, C. Nusbaum, R. M. Myers, M. Brown, W. Li, X. S. Liu, Model-based analysis of ChIP-Seq (MACS). Genome Biol 9, R137 (2008).

74. A. R. Quinlan, I. M. Hall, BEDTools: a flexible suite of utilities for comparing genomic features. Bioinformatics 26, 841–842 (2010).

75. G. Finak, A. McDavid, M. Yajima, J. Deng, V. Gersuk, A. K. Shalek, C. K. Slichter, H. W. Miller, M. J. McElrath, M. Prlic, P. S. Linsley, R. Gottardo, MAST: a flexible statistical framework for assessing transcriptional changes and characterizing heterogeneity in single-cell RNA sequencing data. Genome Biol 16, 278 (2015).

76. A. Dobin, C. A. Davis, F. Schlesinger, J. Drenkow, C. Zaleski, S. Jha, P. Batut, M. Chaisson, T. R. Gingeras, STAR: ultrafast universal RNA-seq aligner. Bioinformatics 29, 15–21 (2013).

77. B. Li, C. N. Dewey, RSEM: accurate transcript quantification from RNA-Seq data with or without a reference genome. BMC Bioinformatics 12, 323 (2011).

78. W. E. Johnson, C. Li, A. Rabinovic, Adjusting batch effects in microarray expression data using empirical Bayes methods. Biostatistics 8, 118–127 (2007).

79. H. Li, R. Durbin, Fast and accurate long-read alignment with Burrows-Wheeler transform. Bioinformatics 26, 589–595 (2010).

80. M. V. Kuleshov, M. R. Jones, A. D. Rouillard, N. F. Fernandez, Q. Duan, Z. Wang, S. Koplev, S. L. Jenkins, K. M. Jagodnik, A. Lachmann, M. G. McDermott, C. D. Monteiro, G. W. Gundersen, A. Ma’ayan, Enrichr: a comprehensive gene set enrichment analysis web server 2016 update. Nucleic Acids Res 44, W90–97 (2016).

81. C. Y. McLean, D. Bristor, M. Hiller, S. L. Clarke, B. T. Schaar, C. B. Lowe, A. M. Wenger, G. Bejerano, GREAT improves functional interpretation of cis-regulatory regions. Nat Biotechnol 28, 495–501 (2010).

82. C. L. Smith, J. T. Eppig, The mammalian phenotype ontology: enabling robust annotation and comparative analysis. Wiley Interdiscip Rev Syst Biol Med 1, 390–399 (2009).

83. Y. Zhou, B. Zhou, L. Pache, M. Chang, A. H. Khodabakhshi, O. Tanaseichuk, C. Benner, S. K. Chanda, Metascape provides a biologist-oriented resource for the analysis of systems-level datasets. Nat Commun 10, 1523 (2019).

